# The combined impact of killer whale consumptive and non-consumptive effects on northern sea otter population viability in the Western Aleutians Archipelago, Alaska

**DOI:** 10.1101/2023.01.30.526353

**Authors:** Tim Tinker, Paul Schuette, Benjamin Weitzman, Joseph Eisaguirre, William Beatty

## Abstract

Predators can alter the abundance, distribution, and behavior of prey populations through consumptive and non-consumptive effects. In the Aleutian Archipelago of Alaska, killer whales (*Orcinus orca*) are considered the most probable cause of northern sea otter (*Enhydra lutris kenyoni*) population declines in the southwestern Alaska stock, which led to their listing as a threatened distinct population segment under the Endangered Species Act. Much of the research attention in the Aleutian Archipelago region has focused on the consumptive effects of killer whales on sea otter population dynamics. Here, we explore non-consumptive effects by accounting for restricted sea otter habitat use within discrete predation refuges characterized by areas of shallow, complex reef habitats close to shore. We constructed Population Viability Analysis (PVA) models that incorporated sea otter count data collected by aerial and skiff-based methods over six decades (1959 – 2021) to inform uplisting (to endangered) and downlisting (delisting from ESA) criteria. Our models incorporated both density-dependent effects and density-independent effects on sex and age structure, which we termed predation hazards. Prior to 1990, predation hazards were negligible, fluctuated at high values between 1990 and the early 2000s, and then declined as sea otter populations reached low densities. We estimated a current regional abundance of 2,405 sea otters (95%CI = 1,734 – 3,238) in the Western Aleutians Management Unit. Our base PVA model that considered only inter-island fragmentation indicated the risk of the regional sea otter population becoming endangered was <5% when there were at least 1,500 otters (95%CI = 1,200 – 2,100), and provided a delisting threshold of 2,100 sea otters. A PVA model that accounted for restricted habitat use of sea otters within discrete predation refuges (i.e. inter-island and intra-island fragmentation) indicated a less encouraging potential for sea otter recovery. The probability of the population becoming endangered increased to >10% and the delisting threshold increased to >10,000 sea otters (nearly 5x higher). Our results indicate sea otters within fragmented predation refuges could be more susceptible to the effects of stochastic processes with potentially limited ability for rescue effects. Overall, our research reveals the importance of evaluating both consumptive and non-consumptive effects when considering conservation and management plans for at-risk populations thought to be limited by a predator.

## Introduction

Predators can influence prey through consumptive effects (i.e., predation) as well as non-consumptive effects such as predator-induced changes in habitat use (Werner et al. 1983), aggregation (Wrona and Dixon 1991), foraging (Lima et al. 1985), or other behaviors intended to limit the risk of predation (Lima and Dill 1990). The total combined effect of consumptive and non-consumptive effects can have profound impacts on prey population dynamics (Creel and Christianson 2008) with concomitant effects on ecosystem processes (Schmitz et al. 1997, Fortin et al. 2005, Hawlena and Schmitz 2010).

The impact of consumptive effects on prey population dynamics and ecosystem processes has been well documented in the Aleutian Archipelago of southwestern Alaska (Estes and Palmisano 1974, Estes et al. 1978, 1998). Following nearly two centuries of overexploitation during the maritime fur trade era (18^th^ – early 20^th^ centuries), northern sea otters (*Enhydra lutris kenyoni*) grew rapidly from remnant populations in the Rat and Andreanof island groups, approaching carrying capacity at some islands (e.g. Amchitka, Adak; Kenyon 1969). The abundance of sea otters at these islands had a density-dependent, limiting effect on herbivorous sea urchin populations through predation, which in turn, influenced the diversity, abundance, and structure of kelp forests as a trophic cascade (Estes and Palmisano 1974, Estes et al. 1978). In the early 20^th^ century, the sea otter population declined drastically again, not by harvest, but likely as a result of predation by killer whales (*Orcinus orca*) (Estes et al. 1998, Tinker et al. 2021a). Although northern sea otters and transient, mammal-eating killer whales have co-occurred for millennia, a potentially small sub-set of killer whales in the North Pacific Ocean was hypothesized to have switched their primary prey preferences or expanded their dietary breadth to include sea otters (Estes et al. 1998, Springer et al. 2003, Williams et al. 2004). Although contentious (Springer et al. 2003, 2008, DeMaster et al. 2006), the killer whale hypothesis was stated as the most probable cause of the sea otter decline when the Southwest Distinct Population Segment (DPS) was listed as threatened under the Endangered Species Act (ESA) in 2005 (U.S. Fish and Wildlife Service 2013), which was then partitioned into five Management Units (MU) - Western Aleutians; Eastern Aleutians; Bristol Bay; South Alaska Peninsula; Kodiak, Kamishak, Alaska Peninsula. Killer whale predation has continued to be the best-supported explanation for the observed sea otter population decline that was rapid (occurred in less than a decade), severe (90% decline from populations near carrying capacity), and both age and sex-independent (Tinker et al. 2021a).

The non-consumptive effects of predators on prey have not been well-studied in nearshore ecosystems of the Aleutian Archipelago. However, sea otters exhibit behavioral patterns in response to killer whale predation indicative of non-consumptive effects. For example, sea otters have adjusted their habitat use and adopted other predator avoidance strategies in response to killer whales, which could potentially have a considerable impact on sea otter population dynamics (Stewart et al. 2015, Tinker et al. 2021a). Remnant sea otter populations at occupied islands are distributed in discrete clusters representative of predation refuges (Stewart et al. 2015). These predation refuges consist of habitats that limit the potential for marine predators to attack from below, and feature shallow, high complexity, reef habitats (e.g “foul” on nautical charts), restricted access points that a killer whale could not swim through, or have escape options to intertidal habitats or on shore. For example, at Adak Island’s Clam Lagoon, sea otters have remained stable over the last several decades despite observed declines elsewhere across the archipelago (Estes et al. 1998). The lagoon’s outlet allows sea otters to pass, but is too shallow for killer whales. Similarly, semi-enclosed bays at Shagak Bay and Bay of Isles have supported sub-populations of sea otters over the past two decades despite declines along adjacent exposed coastlines (Stewart et al. 2015).

Within these predation refuges, sea otters exhibit unique behaviors distinct from their predecessors in the region and from their modern counterparts across the rest of the species’ range (Monson 2021). Specifically, since the observed decline in the 1990s, sea otters are predominantly found in very shallow waters close to shore (<20 m) despite ample food resources further from shore and in deeper waters (Tinker et al. 2021a). These changes in habitat use and resting/foraging behavior may reflect non-consumptive effects through a perceived or realized risk of predation (Gaynor et al. 2019). Sea otter retreat into these predation refuges has allowed low abundant populations to persist for the past two decades, at low but stable densities, but their ability to disperse and repopulate other portions of coastline may be hindered.

Here, we begin to explore how the combination of consumptive and non-consumptive effects by an apex predator, the killer whale, has influenced the spatial and temporal dynamics of northern sea otters in nearshore ecosystems of the Western Aleutians MU. We first explore how predation reduced an abundant sea otter population and fragmented the population into a set of low density island populations. We then consider how non-consumptive effects, evident through the restricted habitat use and altered behavior of otters within each island’s predation refuges, have influenced sea otter populations through time and make projections forward to assess this population’s viability.

We evaluate the impact of consumptive and non-consumptive effects on sea otter population dynamics across space and time through Population Viability Analysis (PVA) (Boyce 1992). We build upon previous analyses that incorporated aerial and skiff-based sea otter population surveys to evaluate the consumptive effects of killer whale predation on the observed sea otter decline and projected forward population dynamics to inform uplisting criteria (from threatened to endangered status) and downlisting criteria (delist from threatened to removal from ESA listing; Species Recovery Plan 2013). In this PVA, we incorporate updated population survey information from 2015 and 2021, which extends the available data to more than six decades (1959 – 2021) of observation. Our model incorporates both density-dependent effects and density-independent effects on sex and age structure, which we term ‘predation hazards’. These predation hazards are allowed to vary across space and time. Overall, our PVA model aims to extend upon the restricted view of killer whale predation as the primary cause and limiter of sea otter recovery, to a more comprehensive view in which we evaluate how both predation and the non-consumptive effects of predation have reduced sea otter populations and hindered their recovery in the Western Aleutians MU.

## Methods

### Overview

We constructed a PVA model for sea otters in the Western Aleutians MU (Figure 1 inset), as a projection matrix that incorporated separate age and sex classes and allowed us to quantify the effects of multiple sources of mortality: 1) age-varying “baseline” hazards and 2) predation hazards as an age-independent process. Our model structure and baseline vital rates were informed by a PVA model described previously in the Southwest Alaska Sea Otter Recovery Plan (U.S. Fish and Wildlife Service 2013) and a more recent analysis of trends in sea otter mortality and abundance in Southwest Alaska (Tinker et al. 2021a).

**Figure 1.**
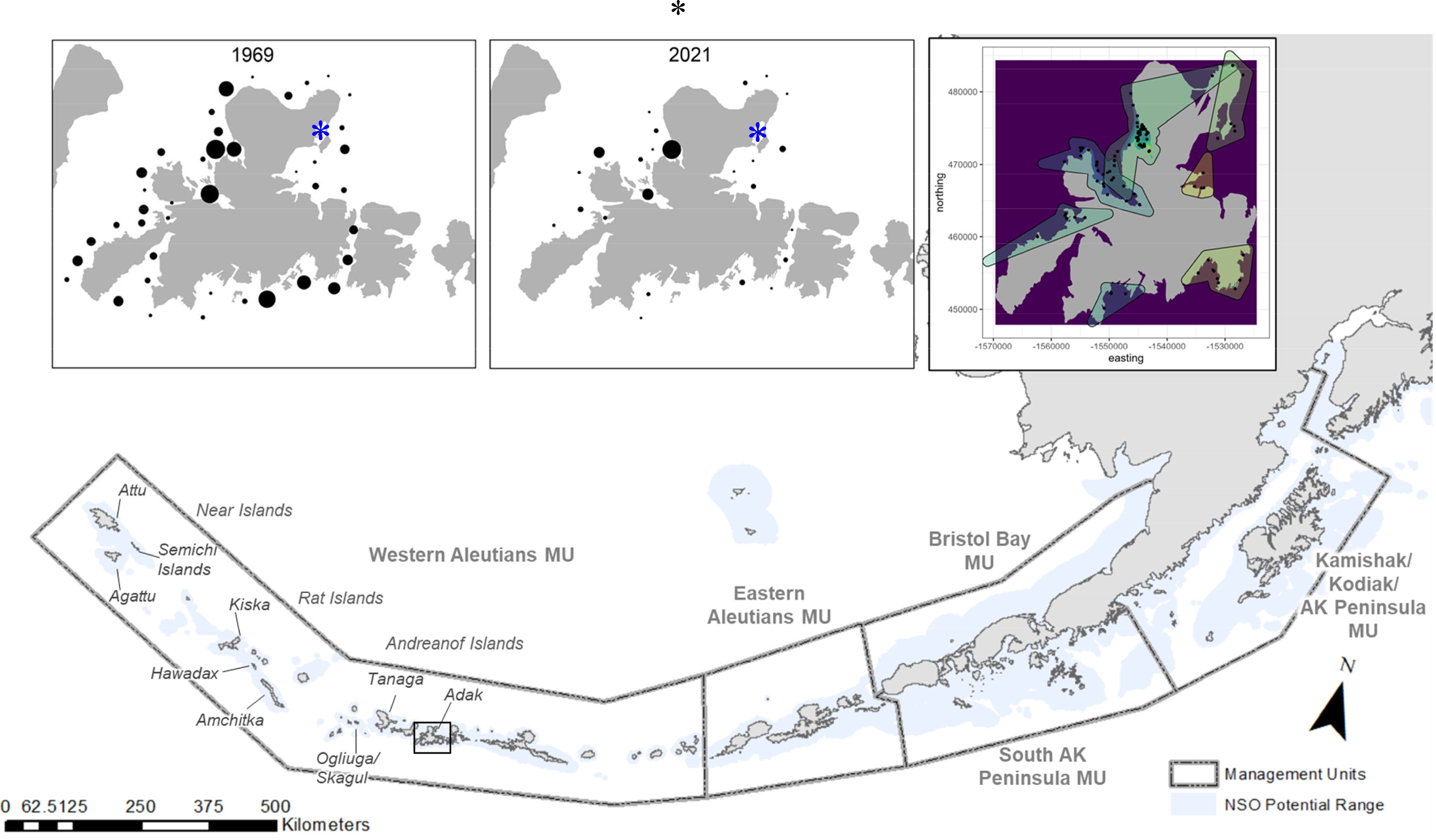
The five Management Units (MU) of the Southwest Distinct Population Segment (DPS) in Alaska. Northern sea otter (NSO) potential range is defined as the 100m isobath within the median ice extent for the month of March from 2001–2020, and restricted to the 24 nautical mile Contiguous Zone maritime boundary. Long-term population monitoring of sea otters spanning six decades in the Western Aleutians MU tracked changes in sea otter population abundance across islands and within islands through time. Top left panel: Sea otter relative counts (otter counts per section of coastline / total count around the island) from an aerial sea otter survey in 1965. Top center panel: Sea otter relative counts (otter counts per section of coastline / total count around the island) from a skiff-based sea otter survey in 2021. We note that skiff-based surveys generally have a higher detection probability than aerial surveys, and thus, the actual difference in sea otter distribution and relative abundance from 1965 to 2021 is likely greater than illustrated. The blue star indicates Clam Lagoon, which was not mentioned in the summary of the 1965 population survey, but is known to be a sea otter hotspot before and after the 1990s decline. Top right panel: Sea otter detections from 2021 are highly clustered with most individuals detected around Shagak Bay and Bay of Isles (northwest portion of the island) and few scattered otters around the remainder of the island’s coastline.

Our approach consisted of three main steps: 1) we developed a process model of sea otter population dynamics that included spatial structure, density-dependence, environmental and demographic stochasticity, baseline mortality (which varies by density, age and sex) and additional hazards associated with predation; 2) we fit the process model to data from aerial and skiff-based surveys at 13 index islands, using Bayesian hierarchical methods, to obtain estimates of key demographic parameters and their dynamics; 3) the model structure and parameter estimates from steps 1 and 2 were then used to conduct forward simulations of sea otter population dynamics across all islands in the Western Aleutians MU over a 25-year period. The simulations were iterated many times, with different initial values for sea otter density, and the results of these simulations were used to determine a) the range of initial abundances for which there is a significant probability that the population will decline to quasi-extinction within 25 years (the upper end of this range is recommended as an appropriate up-listing to endangered threshold), and b) the range of initial abundances for which there is a significant probability that the population will decline to the up-listing threshold within 25 years. The upper end of this range is recommended as an appropriate down-listing threshold, in this case, removing northern sea otters from ESA protections (i.e., delisted).

### Process model

Several previous analyses of sea otter population dynamics in Southwest Alaska have been conducted, and these studies have encompassed a range of complexities in terms of the treatment of demographic processes, from unstructured simple logistic models (Tinker et al. 2021a) to matrix models that include variation in age- and sex-specific vital rates (Monson and DeGange 1995, Monson et al. 2000). While simple unstructured models can be powerful tools for evaluating spatial and temporal trends (Morris and Doak 2002), the inclusion of age and sex structure can be important for PVA models where small population sizes require realistic treatment of demographic stochasticity and Allee effects (Caswell 2001, Gerber and White 2014). We therefore incorporated age and sex structure in our model, but in a simplified format that reduced the number of parameters to be estimated. Specifically, we tracked abundance separately for the first 3 post-weaning year classes (stage 1 = 0.5-1.5y, stage 2 =1.5-2.5y, stage 3 = 2.5-3.5y) and collapsed ages ≥ 3.5y into a 4^th^ adult stage. We consider this appropriate as adult survival and reproductive rates are much less variable than during the first 3 sub-adult years (Monson et al. 2000). We tracked abundance separately for male and female age classes because sex-based differences contribute to demographic stochasticity at low population sizes.

For each of the resulting 8 age-sex classes (female age classes *i* = 1:4 and male age classes *i* = 5:8) we defined annual survival (*S_i_*) and reproductive output (*R*), recognizing that *R* is the product of two separate processes, birth rate and weaning probability (i.e., pup survival to 6 months). Birth rates have been found to be relatively invariant within and across sea otter populations, with adult female sea otters producing a single pup almost every year, while survival of pups from birth to weaning varies dramatically as a function of population status (Monson et al. 2000). Accordingly, we assume a constant birth rate of 1 pup per year for adult females and we consider variation in *R* to represent variation in pup survival.

We used a competing hazards approach for analyzing causes of variation in annual survival rates of pups (*R*) and independent otters (*S_i_*), including effects of age, density, environmental and demographic stochasticity, and the interactions between multiple sources of mortality. Calculation of vital rates in terms of instantaneous hazard rates (*h*) simplified the inclusion of multiple independent mortality sources (because these are additive for instantaneous hazards) and allowed us to evaluate potential covariates in terms of their effects on log-transformed hazards. Thus, for each year *t* and island *j* we define the instantaneous hazard rate for pups as *h_R,j,t_*, which we estimate as a log-linear function

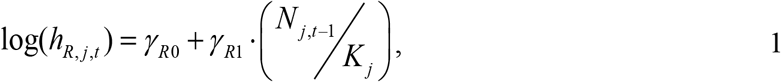

where parameters *γ_R0_* and *γ_R1_* determined the magnitude and density dependent variation in pup survival. We calculated these values from previously published pup survival estimates and provided these values to the model as fixed values (see section “*Baseline survival parameter estimates*”). Equation 1 does not include a term for stochastic effects; however, we estimate and incorporate stochastic effects at a later step to facilitate incorporation of multiple sources of variation, including environmental and demographic stochasticity. The parameter *K_j_* represents the carrying capacity for the *j^th^* population and is calculated as the product of sub-tidal habitat area (*A_j_*, units of km^2^) and island-specific equilibrium density (*Kd_j_*). We induced shrinkage of each *Kd_j_* to a global mean by:

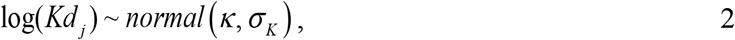

where hyper-parameters *κ* and *σ_κ_* (the mean and standard deviation in log transformed density at *K*) were estimated.

For independent age classes, survival rates are assumed to be affected by two competing sources of mortality: 1) baseline hazards (*h_B_*), which vary by age and sex and represent mortality risks associated with typical density-dependent processes such as nutritional limitation or disease; and 2) predation hazards (*h_P_*), which represent mortality risks associated with predation (and/or other age- and sex-independent mortality factors responsible for the decline). In the case of baseline hazards, we estimate variation in terms of effects on log-hazards:

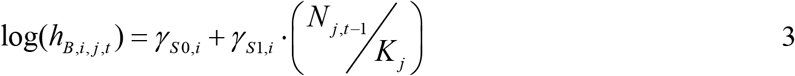

where *γ_S0_* and *γ_S1_* determined the magnitude and density dependent variation in survival and were calculated from previously published survival estimates (see section “*Baseline survival parameter estimates*”) and *K_j_* is the carrying capacity (calculated as for equation 1). In the case of predation hazards, we expected that mortality rates might differ among islands. This could be observed through idiosyncratic features of island habitats, such as the presence of protected or semi-protected lagoons and bays, shallow waters, and other geophysical characteristics that might offer more protection from predation (or less protection in their absence or rarity) as a function of relative population density. For example, in the case of a type-III functional response (Oaten and Murdoch 1975), we might expect predation rates to decline at low densities and with larger amounts of total habitat area (i.e., larger islands with more habitat are more likely to have predation refuges). We modeled these sources of variation in terms of their effects on log-hazards:

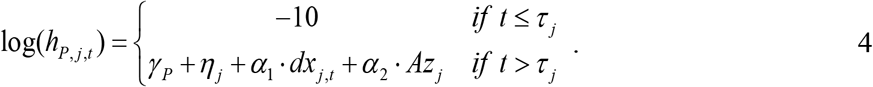

In equation 4, *γ_P_* is the estimated mean log hazard rate from predation; *η_j_* determines island-specific effects and is drawn as a random normal variable with mean of 0 and standard deviation *σ_L_* and *σ_1_* determines the effects of relative density (specified by index *dx*, as defined below); and *α_2_* determines the effects of total habitat area (variable *Az* = habitat area centered and scaled to mean zero and unit variance to facilitate model fitting). We also defined a temporal parameter *τ_j_* as the last year prior to the initiation of the decline at island *j*, and we forced *h_P,j,t_* to a very low value (exp(−10)) for *t* ≤ *τ_j_*, based on published evidence that predation hazards were negligible prior to this time (Estes et al. 1998, Tinker et al. 2021a). The relative density index (*dx*) was formulated to allow for non-linear effects of population density on predation mortality risks (Juliano 2020). For example, per-capita mortality rates could show little variation at high densities but then drop rapidly at low densities when the few remaining animals are concentrated at sites with refuge from predator access (Stewart et al. 2014). Accordingly, we computed *dx* as a non-linear, inverse function of abundance relative to *K*:

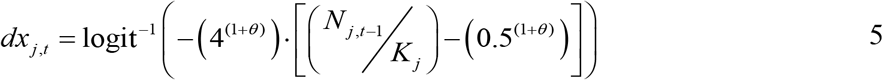

where *θ* is an estimated parameter constrained so that *θ* > 0. Equation 5 results in a 0 – 1 function where *dx* → 0 at high densities (near *K*) and *dx* → 1 at low densities, and the relationship with abundance can take on a range of shapes from approximately linear to strongly sigmoidal (Figure S1).

We accounted for several sources of process error, including environmental stochasticity (*σ_e_*), unexplained variance in predation mortality (*σ_p_*) and demographic stochasticity in the survival of pups and independents. We incorporated each of these sources of variance as additive random terms (normally distributed with mean 0 and sd *σ*) applied to the appropriate log hazard equations. Parameters σ_e_ and σ_p_ were estimated as part of model fitting (see below). To account for demographic stochasticity, we calculated an additional random error term (*σ_d_*) that when added to the log hazard equation produced variance in survival equivalent to sampling from a binomial distribution with the current abundance. Our approach represents an approximation to demographic stochasticity, which we utilized because direct incorporation of demographic stochasticity – treating each survival outcome as a Bernoulli trial – was inconsistent with our treatment of abundance as a continuous latent variable, while the approximation to survival variance inflation simplified incorporation into our instantaneous hazards framework. For a sample of individuals with abundance *n* and expected survival probability *s*, the variance in survival due to demographic stochasticity (binomial sampling error) is estimated as *s*·(1-*s*)/*n*, and annual survival probabilities are calculated from log hazard rates as exp(−exp(log(*h*))). Combining these equations, we used the delta method to transform expected variance in survival to variance in log-hazards:

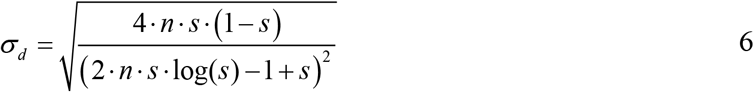

where *n* represents the abundance of pups or independents from a given age/sex class in population *j* at time *t*−1, and *s* represents the expected survival probability given the estimated log hazard rate. Numerical simulations confirmed that variation in log hazards calculated using equation 6 produced a reasonable approximation to binomial sampling variance of survival probabilities even at low abundances (*n*<10).

The process error terms were combined with estimated hazard rates (equations 1, 3 and 4) to calculate annual survival rates for pups and independents:

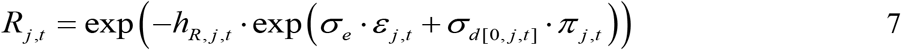

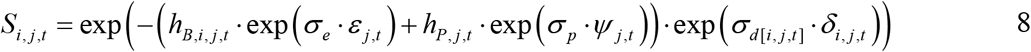

where *ε_j,t_*, *π_j,t_*, *Ψ_j,t_* and *δ_i,j,t_* each represent standard random normal variables with mean = 0 and sd = 1 and *σ* parameters represent estimated variance parameters (note that multiplying a hazard by an exponentiated variance term is mathematically equivalent to adding that variance term to the log hazard equation). Equations 7 and 8 were solved twice, the first time excluding demographic stochasticity terms (*σ*_*d*[0*,j,t*]_ · *π_j,t_* for pups and *σ*_*d*[i*,j,t*]_ · *δ_i,j,t_* for independents) in order to derive the expected survival probabilities (*s*) required to calculate *σ*_*d*[0*,j,t*]_ and *σ*_*d*[*i,j,t*]_ by solving equation 6.

Equations 1 – 8 together describe the estimation of vital rates and their variation over time and space. We combine these vital rates within a stage-structured projection matrix (Caswell 2001) to facilitate computation of annual demographic transitions:

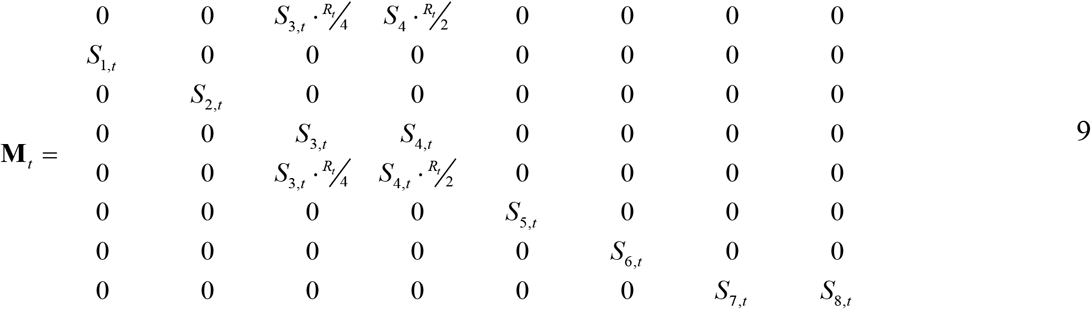

where the elements along the sub-diagonal of columns 1-3 and 5-7 determine survival/growth transitions, diagonal elements in columns 4 and 8 determine survival/persistence of adult females and adult males, and elements in row 1 and row 5 determine reproductive contributions to the juvenile female and juvenile male age classes. We note that reproductive contributions are conditional upon female survival, reflect a 50:50 sex ratio at birth, and stage 3 females are assumed to have a birth rate that is half that of adult females. To simplify notation, we have omitted the “*j*” subscripts, although it is understood that equation 9 is evaluated for each unique island/year combination

To track annual variation in the abundance of each age/sex class (*n_i_*) we used standard matrix multiplication (Caswell 2001):

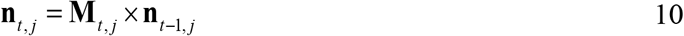

where **n**_*t,j*_ is a vector of stage-specific abundances at year *t*, such that **n**_*t,j*_ = (*n*_1,*t,j*_, *n*_2,*t,j*_,… *n*_8,*t,j*_)’. The sum of vector **n**_*t,j*_ is *N_t,j_*, the total abundance of otters at island j in year *t*. Because equation 10 is recursive, the values of **n**_1,j_ must be estimated: to achieve this we define the initial relative abundance for each island as fitted parameter *F_j_*, representing a fraction of the island-specific equilibrium abundance (*K_j_*). We assume that the initial population would be close to the stationary stage distribution (*ssd*), which can be derived algebraically from **M***t,j*, and we calculate the initial population vector as **n**_1,*j*_ = *F_j_* · *K_j_* · *ssd*.

### Baseline survival parameter estimates

A primary goal of our model was to estimate variation in the consumptive and non-consumptive effects of killer whale predation hazards and their associated impact on sea otter spatio-temporal trends in the Western Aleutians MU. To achieve this goal with the available data, we needed to rely on previously published estimates of baseline survival rates and their density-dependent variation because model identifiability limitations precluded simultaneous estimation of predation hazards and baseline hazard parameters. Fortunately, there are published estimates of age- and sex-specific survival rates and reproductive rates that were measured during tagging studies of low-density and high-density populations of sea otters in the Southwest DPS of Alaska. Estimates of vital rates from the Kodiak Island population (now part of the Kodiak, Kamishak, Alaska Peninsula MU) in the 1980s (Monson and DeGange 1995) correspond to a low density population that was increasing at or near the theoretical maximum rate of growth (*r_max_*~ 0.2), while estimates from Amchitka island in the early 1990s (Monson 1995, Monson et al. 2000) correspond to a stable, high density population that was near carrying capacity, *K*. We note that a population decline had actually begun at Amchitka by 1992; however, the age-at-death distribution on which Amchitka survival estimates were based had accumulated over the preceding decade when the population was still at K (Monson et al. 2000). The published vital rates are reported for annual year classes of males and females (Monson 1995, Monson and DeGange 1995, Monson et al. 2000), and thus required transformation in order to parameterize our stage-based vital rates. We used matrix algebraic methods (Caswell 2001) to calculate stable-stage distributions (*ssd*) associated with each of these sets of vital rates, and then used the age-specific proportions as weights to calculate the weighted mean survival rates and weighted mean weaning rates for adults (>3yrs of age) in each population.

The result of these calculations was a set of estimates for pup survival and independent otter survival for a low-density population (*R_LD_* and *S_LD_*) and a high-density population (*R_HD_* and *S_HD_*) transformed to match our 8 age/sex classes (Available in Supplementary Materials, Table S1). Assuming that linear-interpolation of log-hazard rates between these two extreme cases was a reasonable approach for describing density-dependent variation in survival, we calculated the log hazard parameters for equations 1 and 3 as:

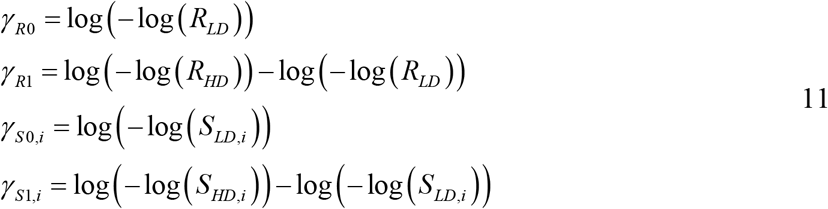

We evaluated this approach by substituting the log hazard values from equation 11 into equations 1 and 3, iteratively evaluating equations 1 – 8 over a range of densities from 0% to 100% of K with *σ*_e_ set to 0.2, calculating the associated projection matrices (equation 9), and then using numerical iterations to calculate expected mean annual growth rates (*λ*) accounting for stochasticity. We note that predation hazards (*h_P_*) were forced to 0 for these calculations, as we were interested in evaluating density-dependent variation in growth rates associated with baseline mortality only. This exercise resulted in a slightly non-linear functional relationship between expected *λ* and relative density (*N*/K), with an average growth rate of 19% per year when *N*/K → 0, and an average growth rate of 0% per year when *N*/K → 1 (Figure S2). The shape of this function was consistent with the expectations of a theta-logistic population growth model with a value of theta between 1 and 2, which is considered reasonable for most marine mammal species (Wade 1998, Taylor et al. 2000).

### Data model

There were two primary data sources available for fitting the process model: aerial-based surveys of all or part of the Aleutian archipelago conducted periodically between 1959 and 2000 (Kenyon 1969, Schneider 1981, Burn et al. 2003, Doroff et al. 2003), and skiff-based surveys for 13 index islands in the Western Aleutians MU (Tinker et al. 2021a) conducted at irregular intervals (but every 3-5 years since ESA listing) at various times between 1970 and 2021 (Table 1, Table S2; Kenner et al. 2021). We utilized the full set of skiff-based data collected from the 13 index islands and the subset of aerial data collected from the same set of islands. We use variable “*V”* to designate survey platform (*V*= 1 for skiff counts and *V*=2 for aerial counts). We used both skiff and aerial data sets for model fitting to maximize statistical power. Relating survey estimates to model-predicted abundance values presented a challenge because counts from the two survey methods are known to be biased low to differing degrees because imperfect detectability, availability (e.g., due to foraging underwater, diving underwater for safety), and observer error. Skiff-based surveys of sea otters in the Aleutians typically have much higher detection probability than aerial-based surveys (Doroff et al. 2003). Based on anecdotal comparisons of skiff counts with near-simultaneous ground-based counts at Amchitka Island (Merritt and Fuller 1977), skiff surveys in the Aleutians are thought to detect between 70 - 90% of all available otters. Unfortunately, we had no quantitative basis for estimating a correction factor for skiff surveys, so we arbitrarily assigned a mean expected probability of detection of 80% for skiff surveys 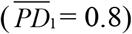. Because many of the islands had multiple instances of near-simultaneous skiff surveys and aerial surveys, it was possible to estimate an average aerial survey detection probability (a pooled estimate for imperfect detectability and availability) relative to 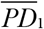 during model fitting, and thus obtain an absolute estimate of mean expected 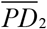 conditional upon our assumed value for 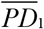. We also allowed for covariates of island size and relative density, based on anecdotal evidence that these factors could influence detection probability:

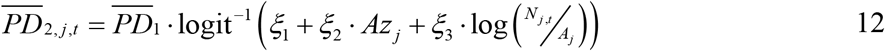

where *Az_j_* is the centered/scaled value of subtidal habitat area (*A_j_*) at island *j*, *N_j,t_* is the estimated true abundance of otters at island *j* at time *t*, and *ξ_1-3_* are parameters to be estimated. We recognized that realized detection probability on any given survey would differ from the expected value due to differing survey conditions, animal distribution, observer experience and sampling error. We incorporated these sources of variation in observer error in terms of a precision parameter (*ϕ*) that we expected would vary by survey platform (*V*) and as a function of sea otter abundance:

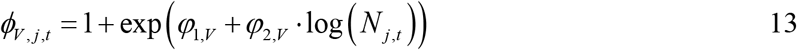

**Table 1.** Summary of meta-data and survey data available for 13 index islands in the Western Aleutians MU.

| Island | Island Group | Habitat<br>Area<br>(km<sup>2</sup>) | Shoreline<br>Length<br>(km) | Longitude | Latitude | Number<br>skiff<br>surveys | Number<br>aerial<br>surveys |
| --- | --- | --- | --- | --- | --- | --- | --- |
| Little Tanaga | Central<br>Aleutians | 72.28765 | 106.39 | -176.1424 | 51.83432 | 3 | 5 |
| Kagalaska | Central<br>Aleutians | 50.59302 | 96.47 | -176.3483 | 51.80512 | 6 | 5 |
| Adak | Central<br>Aleutians | 266.5243 | 372.31 | -176.6274 | 51.78642 | 19 | 5 |
| Ogliuga/Skagul | Central<br>Aleutians | 73.50076 | 48.57 | -178.6497 | 51.60015 | 1 | 4 |
| Kavalga | Central<br>Aleutians | 36.48889 | 29.58 | -178.7934 | 51.5548 | 1 | 4 |
| Semisopchnoi | Rat Islands | 66.21995 | 67.34 | -180.3819 | 51.94937 | 1 | 4 |
| Amchitka | Rat Islands | 305.8711 | 206.71 | -180.9778 | 51.50642 | 11 | 4 |
| Rat | Rat Islands | 70.22691 | 36.17 | -181.697 | 51.79988 | 5 | 4 |
| Kiska | Rat Islands | 221.561 | 196.96 | -182.5307 | 51.97992 | 9 | 4 |
| Buldir | Rat Islands | 31.0936 | 24.73 | -184.0808 | 52.35571 | 1 | 4 |
| Semichis | Near Islands | 164.4185 | 53.21176 | -185.892 | 52.72109 | 9 | 4 |
| Agattu | Near Islands | 165.2205 | 109.76 | -186.4108 | 52.43124 | 7 | 4 |
| Attu | Near Islands | 361.4403 | 260.75 | -187.0699 | 52.89833 | 18 | 4 |

The realized detection probability for a given survey of island *j* at time *t* was estimated as a beta-distributed random variable:

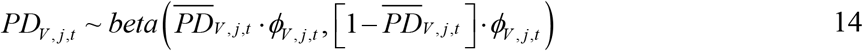

We then related the estimated true abundance (adjusted for detectability) to the observed survey counts (*C*) by assuming the latter were described by a Poisson distribution, thus accounting for additional sampling error in the count data:

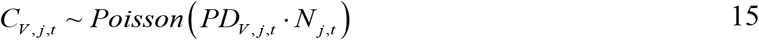

The beta-Poisson data model is not typically used for wildlife survey counts, though it is fairly common in biomedical applications (e.g., Schmidt et al. 2013). For our analysis, the beta-Poisson produced results nearly identical to those obtained using a Negative Binomial distribution to describe survey counts, but with the added benefit of allowing direct estimation of survey-specific detection probabilities.

### Prior model

To ensure that estimates of model parameters are unbiased, and data driven, we utilized vague prior distributions for most parameters (i.e., prior distributions that reflected basic biological or mathematical constraints but did not include information about the specific study system). One exception was the parameter for environmental stochasticity in baseline vital rates, *σ_e_*, the limited survey data prior to the start of the decline precluded reliable estimation of this parameter with a noninformative prior, Accordingly, we set the prior for *σ_e_* as a normal distribution with mean of 0.2 and standard deviation of 0.05, as this distribution described the range of previously reported estimates of environmental stochasticity in log growth rates or survival rates for sea otters (Tinker et al. 2019a, 2019b, 2021a, 2021b). For the remaining variance parameters (*σ_K_*, *σ_L_* and *σ_p_*) we used uninformative half-Cauchy(0, 0.1) priors to achieve shrinkage of the marginal posterior distributions towards 0. We used Cauchy(0, 1) priors for parameters *α*, *ξ*, and *φ*, while for parameter *θ* (which was biologically constrained to a range of values from 0 to 5) we used a gamma(1, 1) prior. For log median equilibrium density (*κ*) we used a normal prior with mean of 2 and standard deviation of 1, as this resulted in a broad distribution of potential equilibrium densities between 0 and 50 otters/km^2^ (a range which encompasses all published equilibrium densities for sea otters). For *γ_P_* we used a normal prior with mean of −2 and standard deviation of 2, as this resulted in a broad and non-informative distribution of predation mortality rates. For parameter *F_j_* we set beta(30,1) priors for islands thought to be at or near *K* in 1960 (Burn et al. 2003) and beta(1,3) priors on the remaining populations.

An additional key source of prior information was the timing of the start of the decline, the point at which predation hazards for sea otters increased rapidly. The initial report of the sea otter decline identified the timing of the start of the decline as the late 1980s or early 1990s (Estes et al. 1998), and a more recent hindcast analysis indicates an average timing of the start of decline as 1990, with some variation among islands (Tinker et al. 2021a). For islands that were included in the trend analysis by Tinker et al. (2021a) we set *τ_j_* to the reported estimated temporal breakpoint for that island. For all remaining islands, we set *τ_j_* to 1989, the average year prior to the initiation of decline (Tinker et al. 2021a).

### Model fitting

The observed data (*C*) constrained the possible values of unknown parameters in the process model and data model, allowing us to approximate marginal posterior distributions for these parameters using standard Markov Chain Monte Carlo (MCMC) methods. We used R (R Core Team 2022) and Stan software (Carpenter et al. 2017) to code and fit the model, saving 10,000 samples after a burn-in of 500 samples. We evaluated model convergence by graphical examination of trace plots from 5 independent chains and by ensuring that Gelman-Rubin convergence diagnostic (r-hat) was ≤1.05 for all fitted model parameters. We conducted posterior predictive checking (PPC) to evaluate model goodness of fit, graphically comparing the distributions of observed vs. out-of-sample count data generated from the same estimated distributions. We also used the sum of the Pearson residuals to compare the fit of observed and out-of-sample data (Gelman et al. 1996), creating scatter plots of the posterior distribution of summed Pearson residuals for each data set. In the case of well-fitting models, points in such a plot should be evenly distributed around a line with slope 1. We then calculated the Bayesian-P value (also using the sum of squared Pearson residuals test statistic), where 0.05< Bayesian-P< 0.95 indicates reasonable model fit (Gelman et al. 1996). We summarized results by reporting the mean, median, 90% and 95% Credible Intervals (CI) of posterior distributions for all model parameters, and by graphically evaluating estimated trends in abundance and predation mortality at select islands.

### Forward simulations

#### Effect of inter-island fragmentation

We used forward simulations of the PVA model to evaluate how current population size could affect future risks of quasi-extinction or endangerment, following methods described in the Species Recovery Plan (U.S. Fish and Wildlife Service 2013). In order to parameterize these forward simulations, we first needed to update our expectation of the distribution of sea otters in the Western Aleutians MU. In the Recovery Plan, the initial distribution for simulations was based on the survey results from the 2000 aerial survey (the last measurement of abundance across the entirety of the Aleutian Archipelago), which is now outdated. Our current model allowed us to update this distribution based on newer survey data and model-estimated dynamics, accounting for continued predation mortality with the effects of density and habitat area incorporated.

Drawing from the joint posterior distribution of the parameters, we iteratively parameterized the process model and projected abundance forward for all islands in the Western Aleutians MU from 2000 to 2021. For random effects we used the estimated posterior distributions for specific effect levels where possible (e.g., for island/year combinations where surveys had occurred) and drew from the appropriate hyper-parameter distributions for unobserved random effect levels. This resulted in posterior distributions of estimated abundance for all islands in 2021 that reflected appropriate levels of uncertainty for each island. We used these estimated abundances (re-scaled to sum to 1) as the probability parameters (Λ_*j*_) for a multinomial distribution used to randomly assigned starting populations at islands for forward simulations.

Forward simulations consisted of repeated iterations of estimated population dynamics over a 25-year period. To initialize a simulation for a specified initial regional abundance, we first distributed animals across islands by drawing from a multinomial distribution with parameters Λ*j*. We used the average stage-distribution from the final year of model fitting to parameterize a second multinomial distribution with which to randomly distribute the initial island populations among stages. Annual dynamics at each island were then calculated following equations 1 - 10, with parameters drawn randomly from the joint posterior distribution, random effects drawn from their appropriate sampling distributions, and all sources of process error (including demographic stochasticity) incorporated as described above for the process model. We allowed for a 10% increase in environmental and predation stochasticity (*σ*_e_ and *σ_P_*) to reflect uncertainty about future conditions, including climate change effects. We conducted two sets of simulations: one to calculate a prospective uplisting threshold (from threatened to endangered), and a second to calculate a prospective down-listing threshold (delisting from the ESA).

To calculate a prospective uplisting threshold, a suite of 10,000 simulations was repeated for 10 different starting densities, ranging from 20 to 1500. For each of the 10 starting densities, we tracked the proportion of simulations that went to quasi-extinction within a 25-year period, a value we designate as Ω. We defined quasi-extinction as the state where there are no remaining islands with at least 5 adult females and at least 1 adult male (U.S. Fish and Wildlife Service 2013). We then fit a nonlinear, log-logistic function to the data set of Ω vs. starting density, using the LL.4 function from the R package “drc” (Ritz et al. 2016), as this functional form resulted in an R^2^ of >0.95. We used the fitted curve to solve for the density at which Ω = 0.05; this value represents the starting density associated with ≥ 5% probability of quasi-extinction within a 25-year period. To account for uncertainty, we used the upper 95% prediction interval from the logistic function to determine the critical density, Ω^05^, which has been suggested as an appropriate value for the uplisting threshold, L^U^. Note that if the estimated Ω^05^ is very low (less than 500), considerations such as maintenance of genetic diversity may be more important than demographic viability, and thus we suggest setting L^U^ to the maximum of 500 or Ω^05^.

To calculate a prospective downlisting threshold, a second suite of 10,000 simulations was repeated for 10 starting densities ranging from 100 to 10,000. For each starting density we recorded ω, the proportion of simulations in which relative abundance drops below L^U^ (the suggested up-listing threshold). Following the same methods described for uplisting simulations we fit a nonlinear logistic function to ω vs starting density, and then solved for the starting density at which ω = 0.05. Above this density there is <5% probability of the population becoming endangered (dropping below L^U^) within 25 years. To account for uncertainty, we use the upper 95% prediction interval from the logistic function to determine the critical density, ω^05^, which is suggested as an appropriate value for the delisting threshold, L^D^.

#### Effect of within-island population fragmentation

The simulations described in the previous section follow the same approach as the original Aleutian sea otter PVA (U.S. Fish and Wildlife Service 2013), whereby individual island populations are treated as single cohesive demographic units. However, recent skiff surveys at index islands in the Western Aleutians MU suggest that sea otters at many islands, especially larger islands, are not distributed continuously, but rather have become fragmented due to predation-driven population declines into discrete spatial clusters, or sub-populations, with extensive stretches of unoccupied shoreline among clusters (Figure 1). We expect this clustered distribution of sea otters could have additional consequences for sea otter population viability if island populations no longer behave as cohesive demographic units but instead represent meta-populations within each island with limited dispersal of animals occurring between sub-populations. To explore the potential consequences of spatial fragmentation within each island for future population viability, we conducted another set of simulations in which island populations were structured as metapopulations.

To design these spatially structured simulations, we first had to specify the relationship between island size (which we characterize in terms of length of coastline, in km) and the number of discrete population clusters, or sub-populations. To achieve this, we used skiff survey data from the 8 index islands surveyed in 2021 whose coastline lengths ranged from 16.5km to 381km. We applied a circular clustering procedure based on von Mises mixtures to the distribution of otters around the coastal perimeter of each island. To implement the clustering, we used cartesian coordinates of individual otters or groups of otters counted on the survey, projected these to the nearest location on the 2-dimensional coastline, and calculated their distance from an arbitrary origin. These distances served as polar coordinates on a circle with circumference equal to the coastline length of each island (calculated from GIS coastline layer at scale 1:24,000 and excluding offshore rocks and islets). The polar coordinates were transformed and projected to cartesian coordinates on the unit circle for clustering. We used the R package “movMF” (hornik and grün 14) to fit a series of von Mises mixtures to the circular otter locations for each island. We fit mixtures with *k* = 1,2,…10 mixture components and chose the value of *k* corresponding to the lowest BIC to be the number of subpopulation clusters for each island. The clustering for Adak Island is plotted as an example (Figure 1c).

We next fit a Bayesian general linear model with number of clusters as the response and island coastline length as the predictor. We determined that the distribution of numbers of clusters were over-dispersed relative to expectations of a Poisson, so we used a negative binomial distribution with log link function. We re-scaled the predictor variable (coastline distance) to units of 100km to improve fitting, and used the “rstanarm” package in R (Goodrich et al 18) to fit the glm, saving the estimated intercept and slope (*β_0_* and *β_1_*) for the linear model of log-transformed mean expected number of clusters vs. coastline distance, as well as the estimated overdispersion parameter (*υ*) from the negative binomial distribution. The posterior samples of these parameters were then used in iterated forward simulations to generate random expected numbers of sub-populations (*k*) for occupied islands.

The methods for conducting forward simulations were generally similar to those described in the previous section, except that matrix-projections were adjusted to incorporate spatial-structure within island populations for all cases where the randomly assigned number of sub-populations *k* >1. Spatial structure was accommodated by several additional algebraic steps. First, we created a block diagonal matrix of demographic matrices (**M**) across the *k* different sub-populations for island *j*:

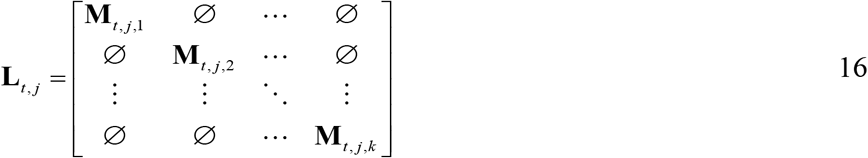

where ∅ represents an 8 by 8 matrix of 0s. We note that each sub-population demographic matrix **M**_*t,j,k*_ is assumed to reflect the same set of underlying vital rates, but will experience differing degrees of demographic stochasticity depending on the number of animals in each sub-population (demographic stochasticity was incorporated according to methods described for the process model, but with the number of animals reflecting the sub-population rather than island population).

To allow for movements of animals between sub-populations within each island, we next defined an inter-population connectivity matrix, **IP**, with dimensions *k* by k:

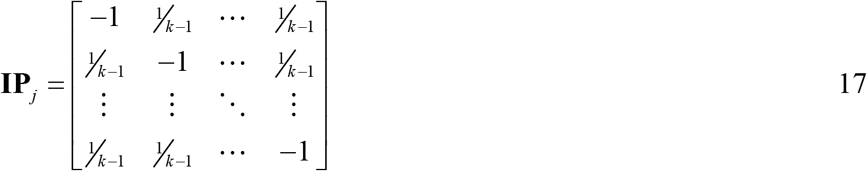

where the values of non-diagonal elements assume that a disperser from sub-population *k_1_* is equally likely to arrive at any other sub-population *k_2_* where *k_2_*≠*k_1_*. (the diagonal was set to −1 to facilitate combining movements and demographic transitions as descried below). We also defined dispersal probability matrix **V**, an 8 by 8 matrix whose cells were set to 0 except for the diagonal, whose values described stage-specific annual dispersal rates (the annual probability that an otter disperses away from its starting sub-population). We set dispersal probabilities to 0.05 for sub-adults and 0.01 for adults of both sexes, based on previously published estimates of sea otter movement rates over similar spatial scales (Garshelis and Garshelis 1984, Ralls et al. 1996, Tinker et al. 2008, Laidre et al. 2009, Tarjan and Tinker 2016, Breed et al. 2017). We then combined matrices **L**, **IP** and **V** to calculate annual dispersal and demographic transitions in a single step:

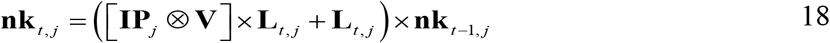

where ⊗ represents a Kronecker product and **nk**_*t,j*_ is a meta-population vector of length 8 × *k_j_* (the number of sub-populations at island *j*), and tracks the number of individuals in each stage, in each sub-population: **nk**_*t,j*_ = (*n*_1,1,*t,j*_, *n*_2,1,*t,j*_,… *n*_8,1,*t,j*_, *n*_1,2,*t,j*_, … *n*_8,k,*t,j*_)’. We initiated **nk**_*t,j*_ at *t*=1 by randomly distributing the starting population for island *j* among the *k_j_* sub-populations using a multinomial distribution (assuming equal likelihood of occupancy for all sub-populations).

We then compared the results of simulations having within-island population fragmentation to the baseline simulation results where island populations are assumed to be homogenous. Specifically, we evaluated how fragmentation within islands would be expected to affect the recommended de-listing threshold compared to if we only considered connectivity among island populations.

## Results

### Model fitting

The process model was fit successfully to the survey data, with well-mixed chains and convergence for all parameters (r-hat < 1.05; Table 2). Goodness of fit diagnostic tests indicated no apparent lack of fit, with close agreement between observed and out-of-sample observations (Figure S3) and Bayesian-*P* = 0.40. Summary statistics for posterior distributions of all parameters are provided in Table 2.

**Table 2.**
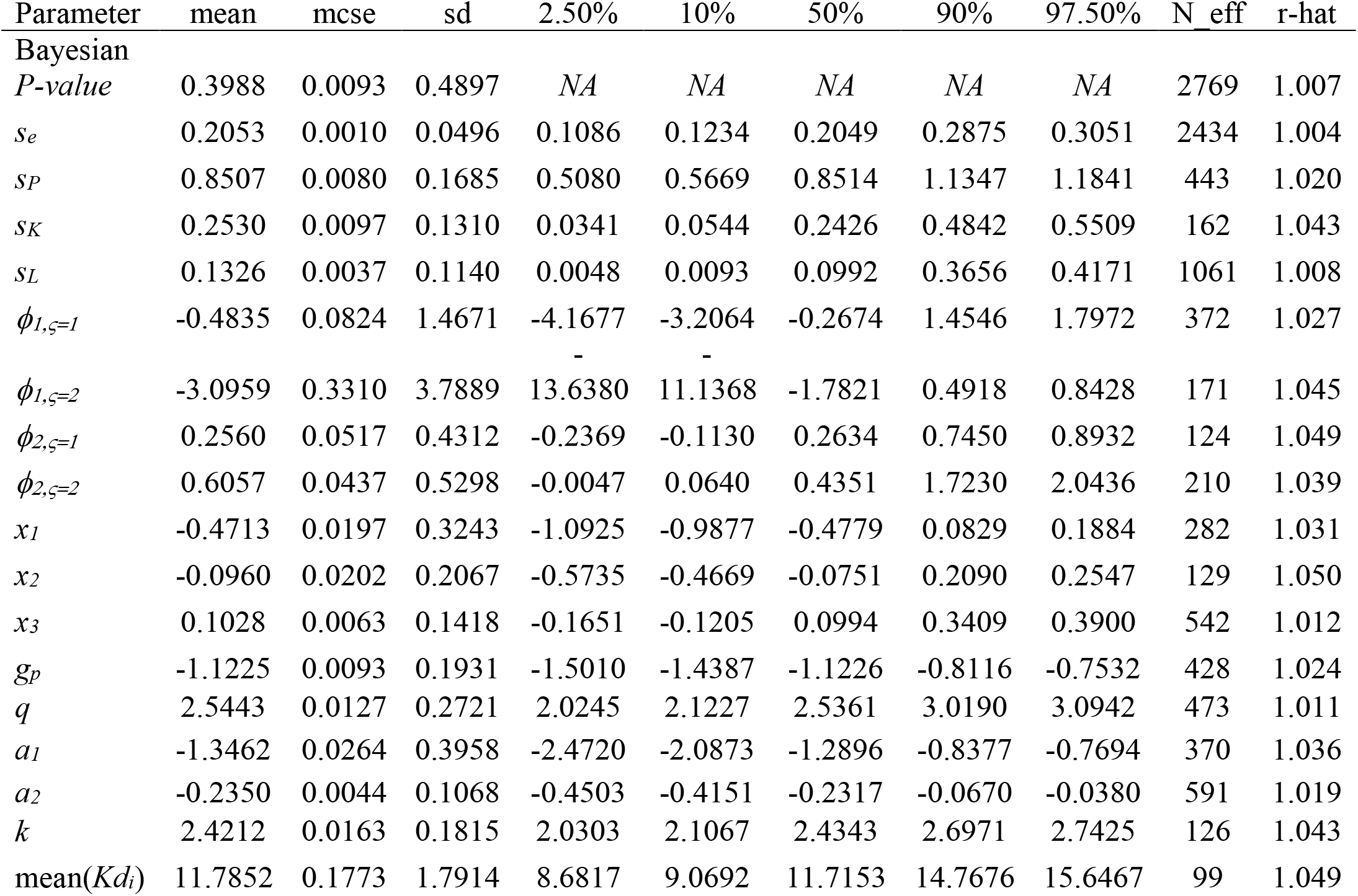
Summary statistics for estimated parameters in a model of sea otter population dynamics for the Western Aleutians MU. Statistics reported for the posterior distributions of each parameter include: arithmetic mean, Monte Carlo standard error (mcse), standard deviation (sd), select quantiles, effective population size (N_eff), and the Gelman-Rubin convergence diagnostic (r-hat).

Key estimated parameters include equilibrium density (*Kd*) and its variability across islands (*σ_K_*), environmental stochasticity in vital rates (*σ_e_*), predation log-hazard rate post-1990 (*γ_P_*), spatial differences in predation hazards (*α_2_* and *σ_L_*), and temporal variation in predation hazards (*α_1_* and *σ_p_*). The mean estimated equilibrium density across the 13 index islands was 11.8 otters/km^2^, although there was variation across islands (SD in log-density, *σ_K_* = 0.26), with *Kd_i_* estimates ranging from 9 to 16 otters/km^2^ (Figure S4). The predation hazard rate after 1990 varied across islands, in part based on differences in island size (larger islands had lower predation hazards, as determined by estimated *α_2_*< 0; Table 2) and in part due to unexplained random effects (*σ_L_* = 0.13). Temporal variation in predation hazards was explained in part by density effects. Specifically, the estimated values of *θ* and *α_1_* parameters (Table 2) resulted in a strongly non-linear relationship between the magnitude of predation hazards and density, with a substantial reduction in predation mortality occurring at low densities (Figure 2). In addition to density effects there were substantial unexplained fluctuations in predation hazards, and the magnitude of this variation in predation hazards (*σ_p_*=0.85) was substantially greater than baseline environmental stochasticity (Table 2).

**Figure 2.**
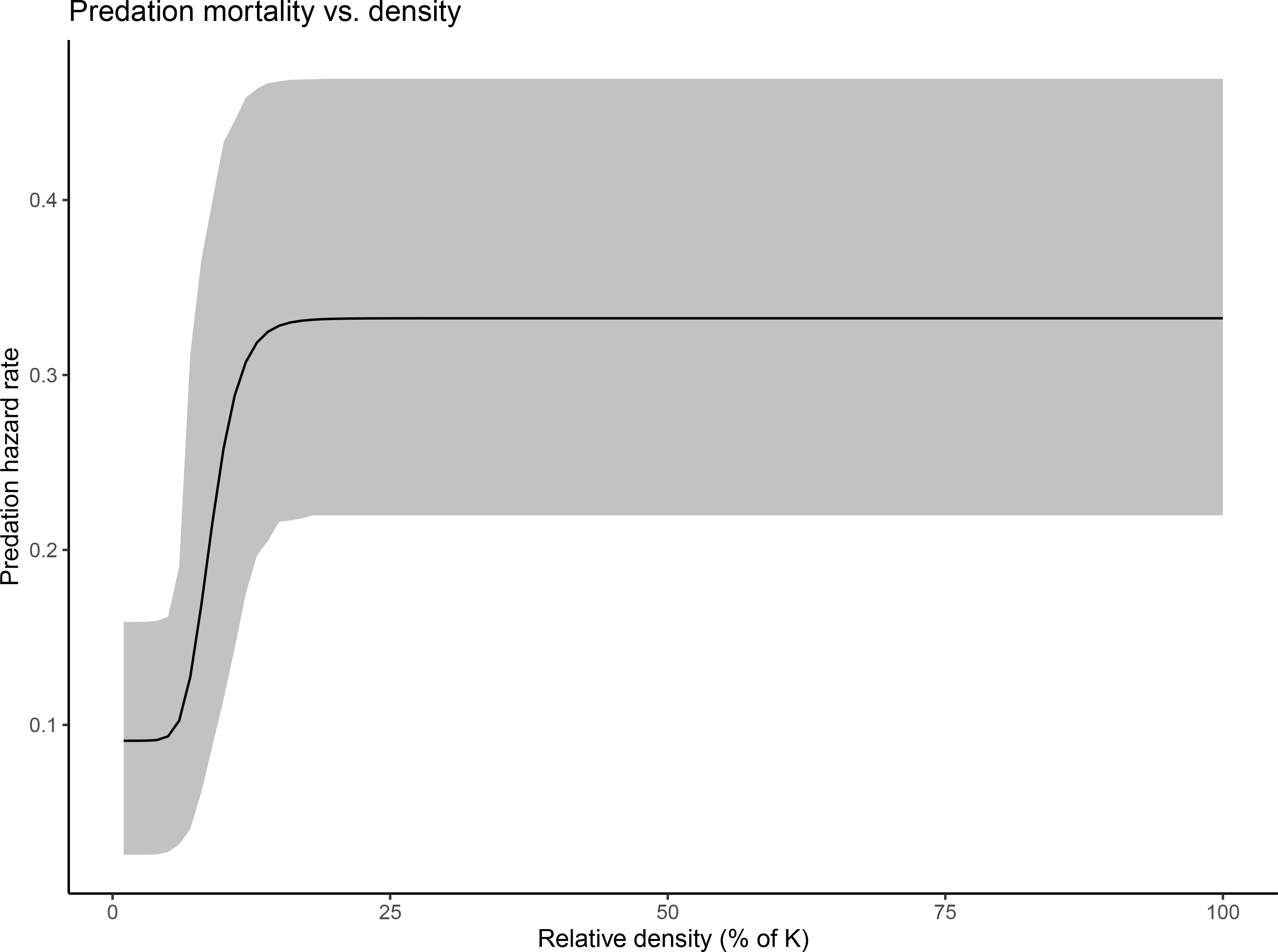
Model estimated functional relationship between the relative density of an island population (as % of carrying capacity, or *K*) and the instantaneous mortality (hazard) rate from predation, averaged across islands and years after 1995.

At a course scale there was a consistent temporal pattern of island-specific predation hazard rates: prior to 1990 predation hazards were negligible, before spiking to high values between 1990 and 1995, fluctuating at high values for about a decade, and then declining over time as local populations reached low densities (Figure 3, a-c). This variation in predation hazards had clear impacts on abundance trends. Prior to 1990, when predation hazards were negligible, population trends varied among islands with some island populations (e.g., Adak, Amchitka in the Andreanof Islands) having reached equilibrium densities by the 1960s while other islands (e.g., Attu in the Near Islands) were still increasing. However, after the spike in predation mortality, all islands declined at consistently high rates (Figure 3, d-f). As densities reached low values, the combination of reduced natural mortality (associated with typical density-dependence) and reduced predation hazards (Figure 2) resulted in a stabilization of trends at densities approximately 5-10% of their pre-decline values (Figure 3, d-f).

**Figure 3.**
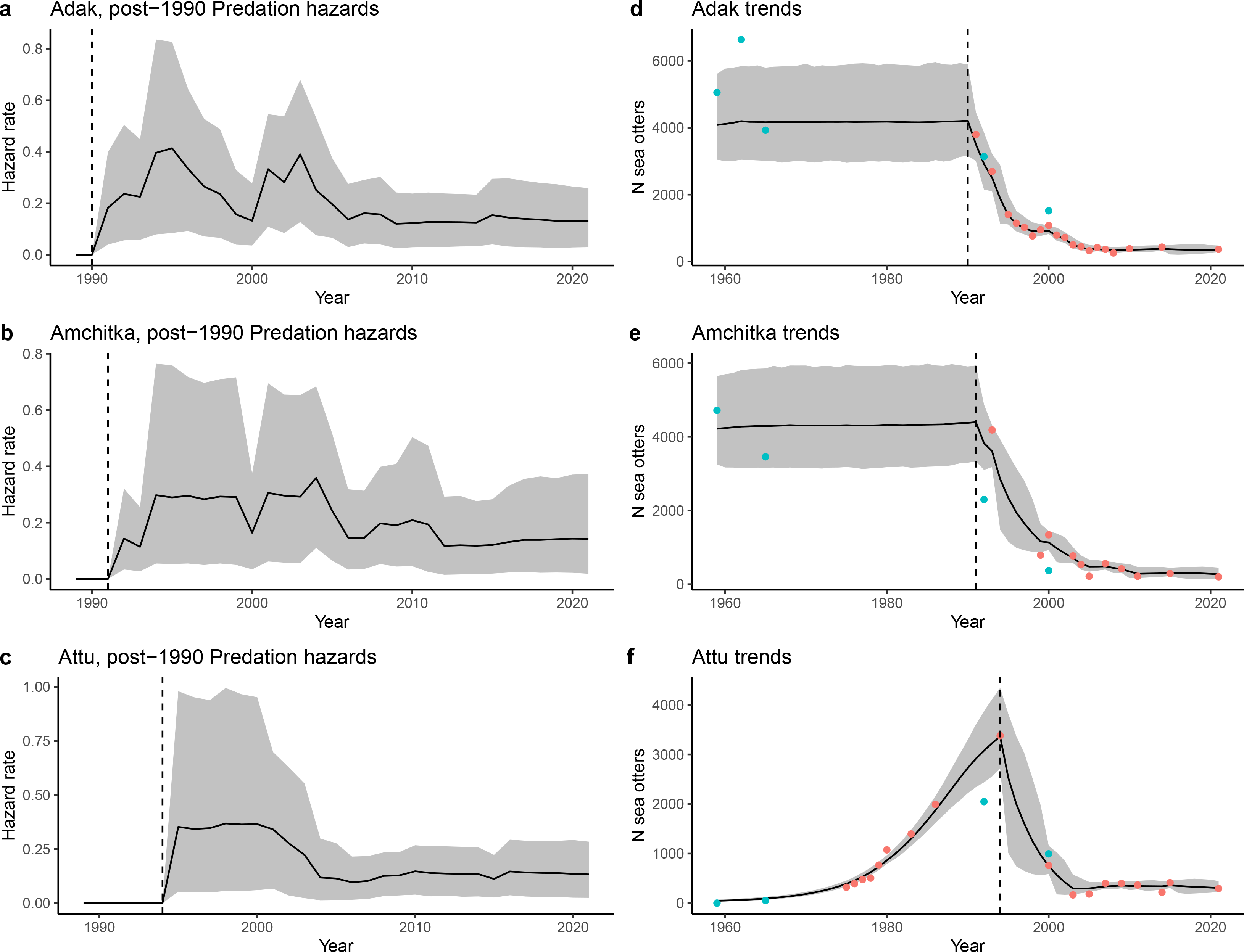
Trends in predation hazard rates and sea otter abundance for 3 index islands in the Western Aleutians MU, each representing one of the 3 main sub-regions: Adak (Adreanof Islands), Amchitka (Rat Islands) and Attu (Near Islands). Panels a-c show the temporal variation in the estimated instantaneous hazard rate from predation, while panels d-f show the temporal variation in abundance. In all cases the dashed vertical line indicates the estimated initiation of predation mortality at the island, the solid line shows the mean estimated value of the plotted statistic and the grey shaded bands indicate the 95% CI of the plotted statistic. For panels d-f the dots represent survey counts (corrected for mean estimated survey detection probability, for consistency with estimated trends): blue dots indicate aerial survey counts and red dots are skiff survey counts.

### Forward simulations

#### Effect of inter-island fragmentation

Each set of model projections for any given island resulted in a distribution of outcomes, with some trajectories going to extinction and some remaining approximately stable or increasing over a 25-year period. In general, larger islands tended to support higher initial populations and thus fewer simulated trajectories declined to quasi-extinction than at smaller islands (Figure 4). We used model simulations to project forward the abundance of all islands from their estimated abundance in 2000 to their estimated abundance in 2021, resulting in a mean estimated regional abundance of 2,405 otters (95%CI = 1,734 – 3,238; Figure 5a) in the Western Aleutians MU.

**Figure 4.**
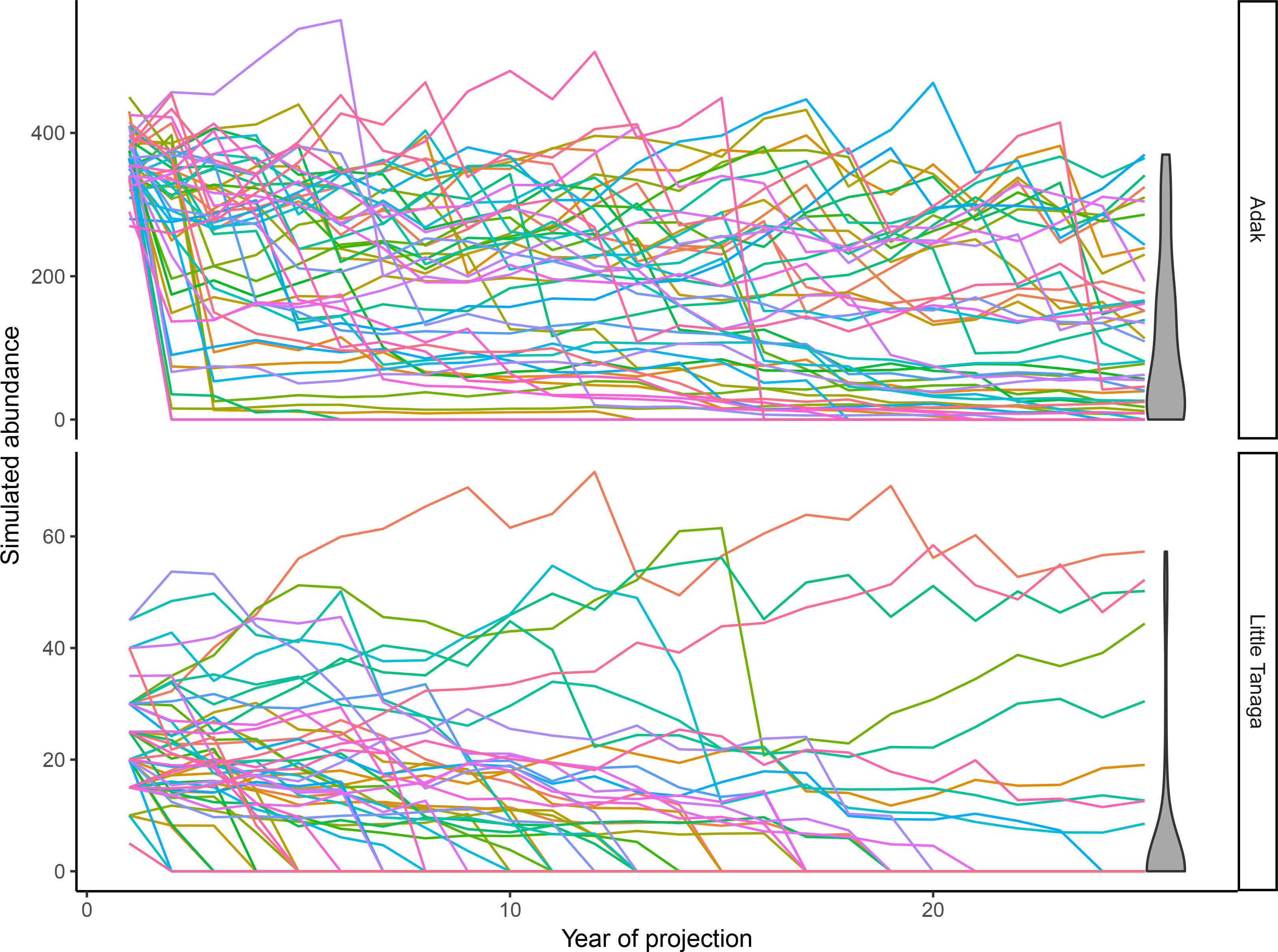
Sample trajectories from iterated simulations of projected future abundance of sea otters at two representative islands in the Western Aleutians MU, Adak (top) and Little Tanaga (bottom). Each of the plotted lines shows the estimated abundance over time for a single simulation, with differences across replicates reflecting parameter uncertainty as well as the effects of process error (environmental stochasticity, predation hazard stochasticity, and demographic stochasticity). The violin plots at right show the distribution of expected abundance at the end of the 25-year projection period. Because Little Tanaga is smaller (106 km coast) than Adak (372 km) and has less available habitat (72 vs. 267 km^2^), there is less potential for refuge habitat and the starting population size is also lower, and a thus a greater proportion of simulated trajectories drop to 0 (quasi-extinction).

**Figure 5.**
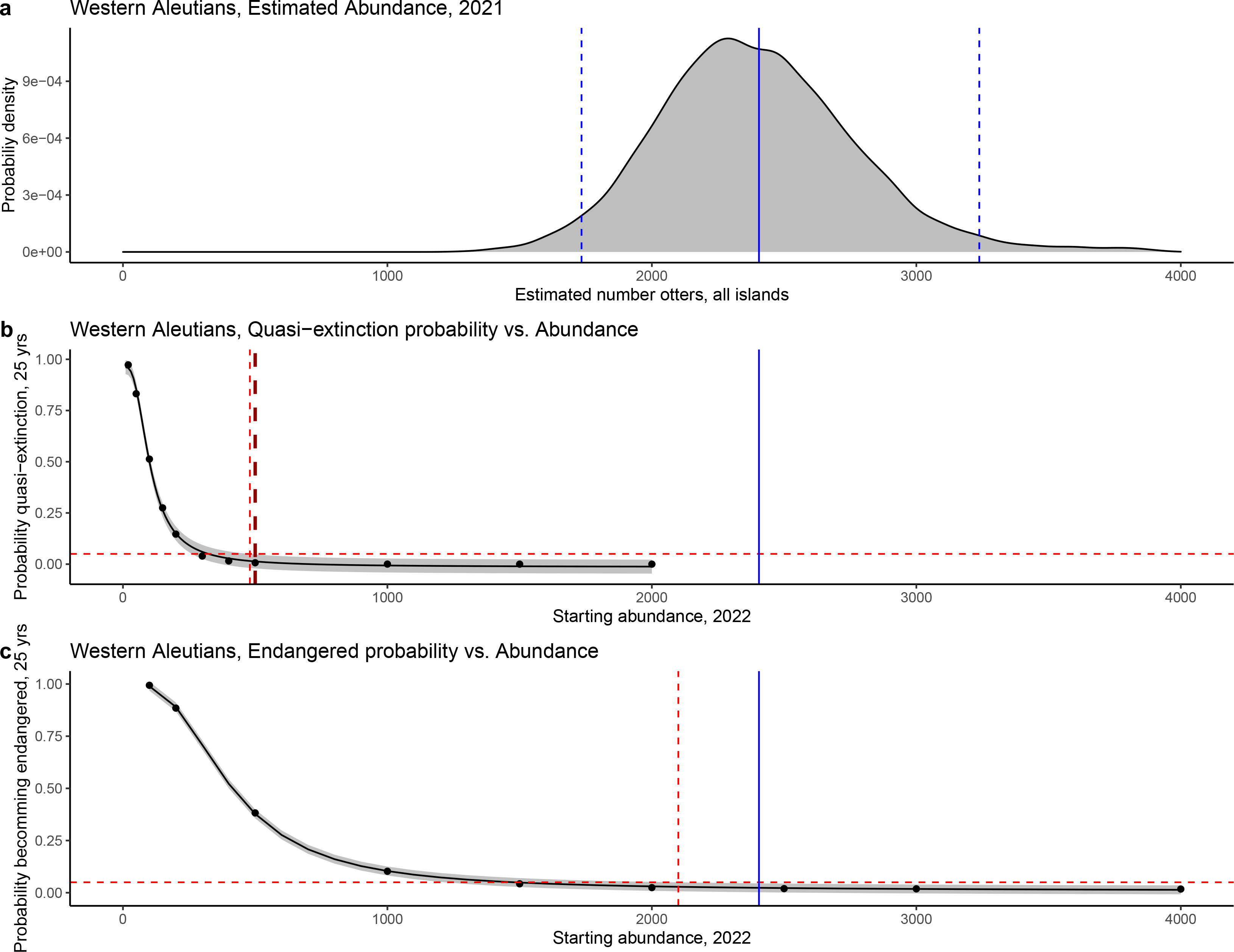
Key results from a population model fit to survey data for sea otters in the Western Aleutians MU, as well as projections of future abundance based on the model. A) a density plot showing the posterior distribution of the estimated total abundance across all islands as of 2021. The blue vertical line indicates the mean estimated abundance and dashed vertical blue lines indicate the upper and lower bounds of the 95% CI. B) The estimated probability of quasi-extinction (QE, defined as no Islands with >5 adult females and >1 adult male) after 25 years, plotted as a function of the total abundance at the start of the 25-year period. A logistic smoother (black line) was fit to simulation results (black dots), with the grey band indicating the 95% prediction interval for the function. The red dashed vertical line indicates “Ω^05^”, the value along the x-axis at which the upper bound of the prediction interval intersects the 0.05 probability value on the y-axis: at abundances >Ω^05^ there is >95% probability that abundance will not drop to QE within 25 years. C) The estimated probability that total abundance will drop below Ω^05^ within 25 years, plotted as a function of the total abundance at the start of the 25-year period. A logistic smoother (black line) was fit to simulation results (black dots), with the grey band indicating the 95% prediction interval for the function. The red dashed vertical line indicates the value along the x-axis at which the upper bound of the prediction interval intersects the 0.05 probability value on the y-axis: at abundances higher than this value there is >95% probability that the total population will not drop below Ω^05^ within 25 years.

We next replicated forward simulations for all islands 10,000 times over a range of initial population sizes, to determine the proportion of simulations going to regional quasi-extinction as a function of regional abundance. This quasi-extinction probability varied predictably as a function of the initial population density, a pattern that was used to determine critical thresholds for extinction or endangerment probability (Figure 5b). The estimated risk of extinction (Ω) was >5% when there were fewer than 320 independent otters, with 95%CI = 260 – 480. Because the upper 95%CI for Ω^05^ was less than 500 animals (Figure 5b), it is suggested that the uplisting threshold (*L^U^*) (from threatened to endangered) be set to 500 for the Western Aleutians MU. Likewise, the risk of the regional population dropping below 500 and thus becoming endangered (ω) was <5% when there were at least 1,500 otters, with 95%CI = 1,200 – 2,100 (Figure 5c). The upper 95%CI for ω^05^ provides a suggested delisting threshold from the ESA for the Western Aleutians MU of 2,100 otters.

#### Effect of within-island population fragmentation

Forward simulations that incorporated within-island fragmentation of sea otter populations into semi-discrete sub-populations in predation refuges provided an additional assessment of the risks of endangerment and local extinction risk. Qualitatively, the results of within-island fragmentation simulations were similar to results with only inter-island fragmentation, with the probability of regional decline to some critical threshold being described by a declining, non-linear function. However, the average probability of the regional population becoming endangered was proportionally higher for each initial population size due to increased effects of demographic stochasticity for very small sub-populations and minimal rescue effect due to limited dispersal between sub-populations (Figure 6). The probability that the population would become endangered was >10% at a population size of 2,100 otters (Figure 6). Under this scenario that includes within-island fragmentation, the upper 95%CI for ω^05^ provides a suggested down-listing (delisting) threshold of >10,000 sea otters, far higher than for simulations without fragmentation (Figure 6).

**Figure 6.**
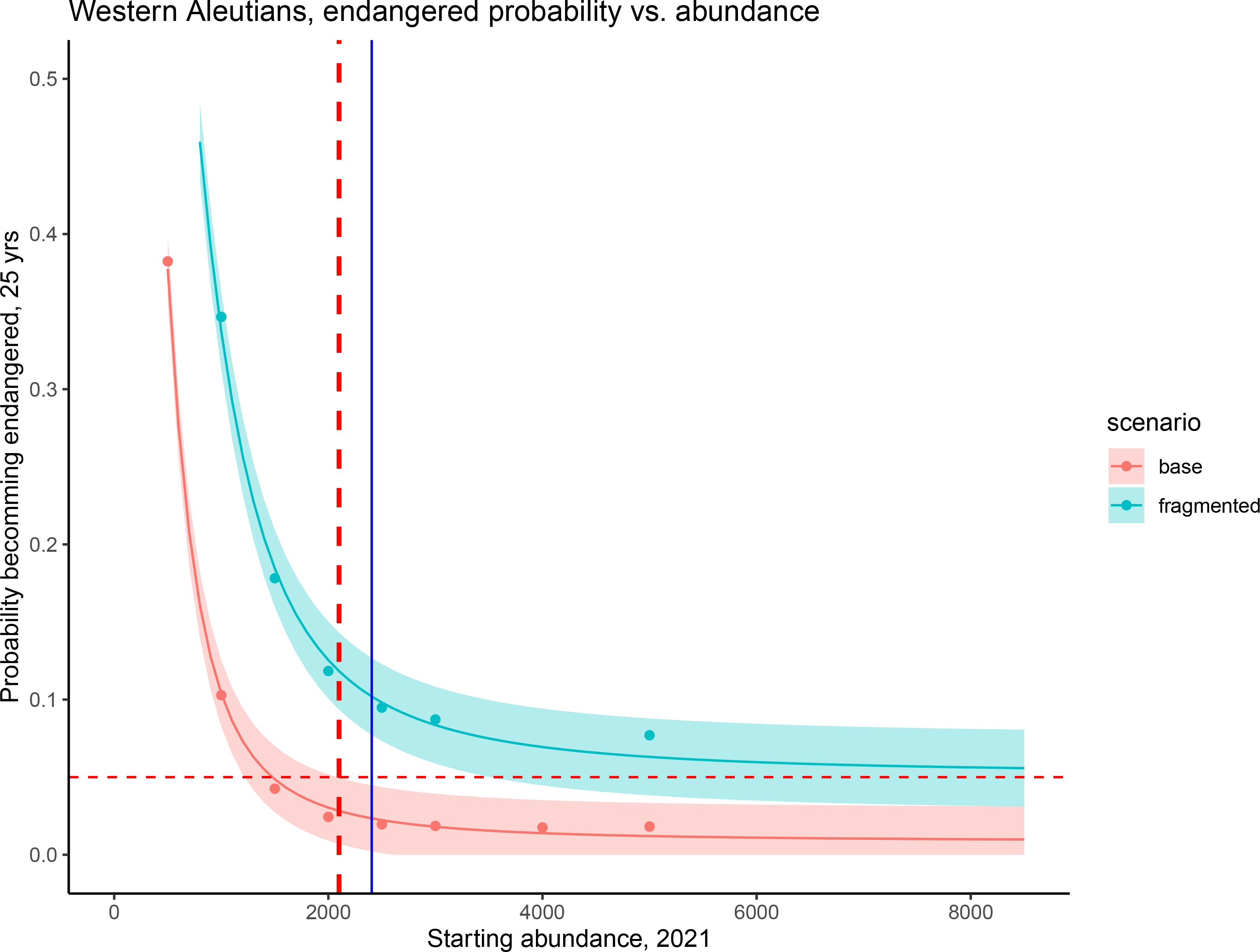
Comparison of projection results from a sea otter population model under two scenarios of sea otter distribution and spatial connectivity. Under the base scenario, each island population is modeled as a single, cohesive population unit. Under the fragmented scenario, sea otters at each island are divided among one or more semi-discreet sub-populations (the number of sub-populations is a stochastic, increasing function of island size), linked by low levels of dispersal. The vertical axis of the plot is the estimated probability that the population will become endangered (total abundance will drop below Ω^05^, the recommended uplisting threshold) within 25 years, while the horizontal axis is the starting abundance of population projection simulations. The functional relationships between endangerment and initial population size under the two scenarios are indicated by the solid lines (mean values of the logistic function) and shaded bands (95% prediction intervals of the logistic functions). The smaller population sub-units in the fragmented scenario are more impacted by demographic stochasticity at low population size, and thus the risk of endangerment at a given abundance is higher for this scenario.

## Discussion

Long term monitoring in the Western Aleutians MU has documented the rise and fall of the sea otter population over the past six decades. The drastic population declines in the 1990s through the early 2000s culminated in the listing of the Southwest DPS as threatened under the ESA in 2005. The cause of the population decline and the inability of this DPS to recover thus far has focused primarily on the consumptive effects of killer whales on sea otters and associated Allee effects from a small, fragmented sub-population (U.S. Fish and Wildlife Service 2013). For the first time, we have updated our view of killer whale – sea otter dynamics to undertake a more comprehensive view of killer whale predation hazards, incorporating both the consumptive effects of predation and the non-consumptive effects of restricted habitat use within predation refuges. Our analyses revealed the combination of consumptive and non-consumptive effects may have a potentially larger impact on sea otter population status and recovery than originally anticipated.

Our initial PVA model focused on inter-island fragmentation (the base model) with sea otter population dynamics driven by the consumptive effects of killer whales. This model provided the first evidence for density-dependent reduction in the age-independent mortality (predation hazards) driving the sea otter population decline. We observed a non-linear relationship between the magnitude of predation hazards and sea otter density, with a substantial reduction in predation mortality occurring at low densities. These results support the notion that a declining sea otter population has been able to persist within predation refuges, especially at larger islands with more complex habitat. Meanwhile, sea otter populations at some small islands with little or no potential predation refuges, such as Semisopochnoi and Ogliuga/Skagul, supported otters in the pre-decline era but currently have no otters. The incorporation of density-dependent variation in predation hazards helps explain the apparent stabilization of trends (at some index islands) at low densities.

The stabilization of population trends in the base model initially suggested improved sea otter prognosis: lower vulnerability to extinction and endangerment - and thus a lowering of the suggested listing thresholds - as compared to previous analyses (U.S. Fish and Wildlife Service 2013). The base model provided a current population estimate of 2,405 otters (95%CI = 1,734 – 3,238) across the Western Aleutians MU, and an uplisting threshold (from threatened to endangered) and downlisting threshold (delisting) of <500 otters and >2,100 otters, respectively. If these estimates were to be used for downlisting and delisting criteria, we could infer the Western Aleutians MU sea otter population is currently straddling the line between threatened status and delisting thresholds.

However, when accounting for the combined effects of consumptive and non-consumptive effects by accounting for sea otter population fragmentation across islands and within-island predation refuges, we show that the potential for sea otter recovery could actually be lower than suggested by earlier analyses (U.S. Fish and Wildlife Service 2013). The PVA model that included both inter-island and within-island dynamics increased the probability of the population becoming endangered (at a starting abundance of 2,100 otters) from <5% probability in the inter-island model (base model) to >10% in the model with inter-island and within-island fragmentation (Figure 6). Likewise, the threshold at which the sea otter population had <5% chance of becoming endangered increased the delisting threshold from 2,100 otters determined by the base, inter-island model to >10,000 otters (nearly 5x higher) in the model with inter-island and within-island fragmentation (Figure 6). If the predation refuge clusters are demographically discrete, our simulations suggest baseline estimates of vulnerability may be optimistic. These smaller sub-populations are more susceptible to effects of demographic stochasticity and, depending on degree of dispersal between discrete clusters, there may be limited potential for “rescue effect” if some clusters become extinct. Thus, results of this version of the model, suggest the current estimate of sea otters (2,100) is vulnerable to stochastic processes and further reductions in the future.

It is important to note that even with our updated uplisting and downlisting thresholds, the Western Aleutians MU is just 1 of 5 MUs in the Southwest DPS. ESA listing status for the entire Southwest DPS is determined based on the collective status of the five MUs (U.S. Fish and Wildlife Service 2013). The Recovery Plan dictates that at least 3 of the 5 MUs must meet recovery criteria in order for delisting to be considered (U.S. Fish and Wildlife Service 2013). These criteria stipulate that in order to be considered for delisting, each MU must 1) achieve and maintain a self-sustaining population, 2) maintain enough sea otters to ensure that they are playing a functional role in their nearshore ecosystem, and 3) mitigate threats sufficiently to ensure persistence of sea otters (U.S. Fish and Wildlife Service 2013). For the Western Aleutians MU, recent analyses indicate this MU is low and stable (Tinker et al. 2021a). However, our current results suggest that the potential for recovery depends on whether the combination of inter-island and within-island fragmentation driven by predation and non-consumptive effects are accounted for. In our simulations accounting for within-island population fragmentation, which we believe reflect a more comprehensive view of predation hazards, the Western Aleutians MU sea otter population is vulnerable to endangerment and shows little sign of growth towards delisting, thus not meeting the first criterion.

For delisting, the Recovery Plan stipulates sea otters must recover to a point at which they are ecologically functional in nearshore ecosystems, i.e., abundant ‘enough’ to consume sea urchins to the extent at which kelp forests can recover (Estes et al. 2010, Estes and Palmisano 1974, Estes et al. 1978). The original research illustrating the ecological functional role of sea otters occurred at islands where sea otters were at or near equilibrium density (e.g., Amchitka), which was estimated to be 15 otters/km^2^ (Burn et al. 2003). Our PVA models provided an updated estimate of sea otter equilibrium density of 11.8 otters/km^2^ (range: 9 – 16 otters/km^2^), which is >3 otters/km^2^ less than the previous estimate. This new estimate of equilibrium density has potential implications for how we view the ecological functional role of sea otters in nearshore ecosystems, and at what density they will be able to restore their ecological functional role in this nearshore ecosystem (Estes and Palmisano 1974, Estes et al. 1978). Prior research linking sea otter densities to nearshore ecosystem state in the Aleutian Archipelago has indicated sea otters can restore ecological function at approximately 6.3 otters/km of coastline, a linear density metric corresponding to approximately 1/3 to 1/2 of carrying capacity (Estes et al. 2010). However, those previous analyses were premised on the assumption that sea otters would exert their top-down effects around the entire coastlines of the islands at which they occurred (as they did prior to the decline); without restriction of movements or space use. This assumption is challenged by our current analyses, which indicate this relative density threshold (i.e., otters/km rather than otters/km^2^) depends not just on the amount of available coastline (i.e., island size), but on the ability of sea otters to access and occupy the entirety of island coastlines. Given that sea otters in the post-decline era are patchily distributed around islands in predation refuges, we expect sea otters to be ecologically functional only within these predation refuges and unlikely along coastlines. Building on observations of sea otter effects in remnant populations, showing that sea otters only influence the habitats where they reside and persist (Stewart et al. 2015), it is likely that sea otters are even more ecologically restricted in where they exert predatory control of benthic grazers, such as sea urchins, and where they can influence nearshore ecosystem state. Further, long-term benthic monitoring using scuba-based methods indicates even with predation refuges, ecological function may be occurring only in shallow waters (<10 ft) very close to shore (<20 m) (Figure 7). Future research could identify the extent to which sea otters are being ecologically restricted and how their functionality varies within predation refuges relative to adjacent coastlines.

**Figure 7.**
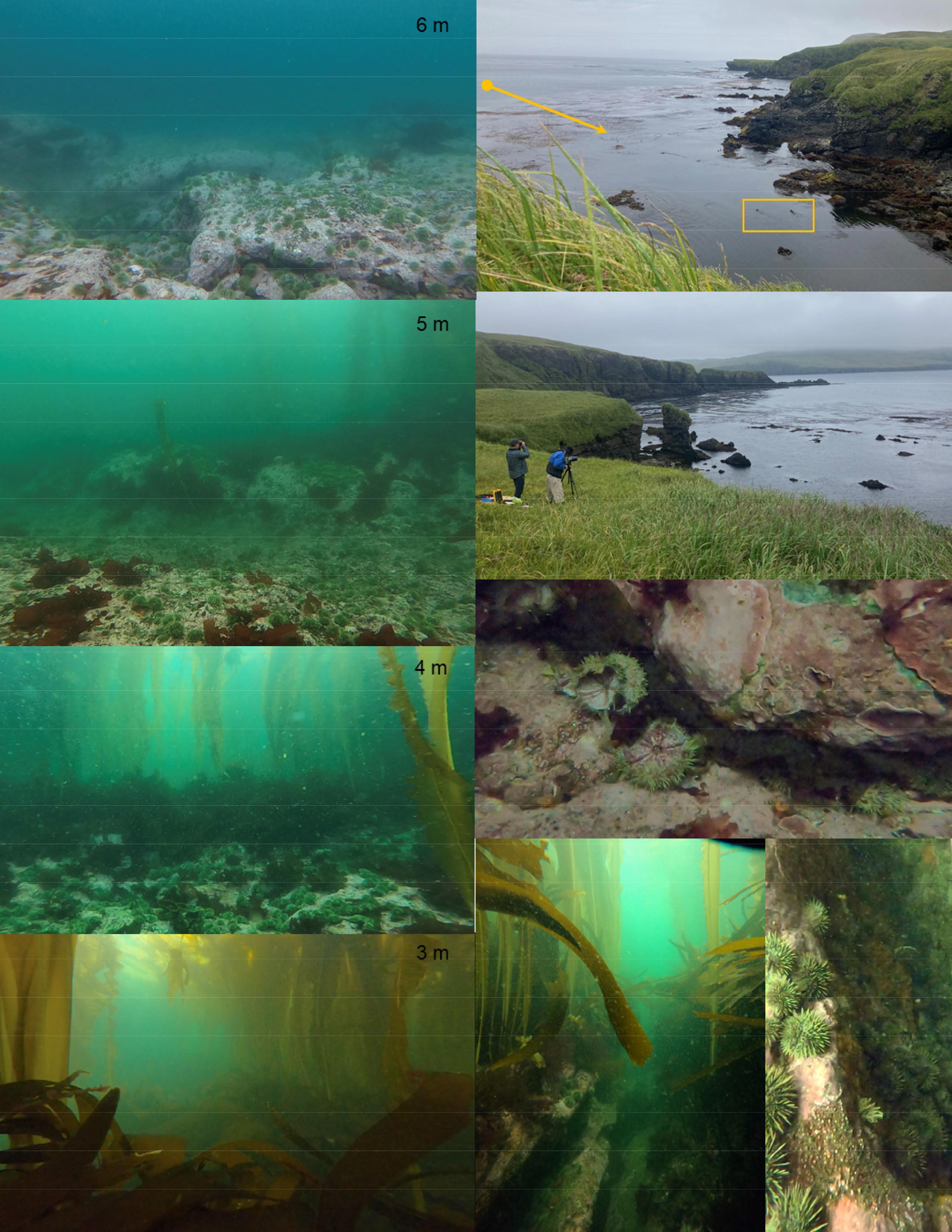
Sea otters can drive nearshore ecosystem state, via predatory control of grazing sea urchins, allowing macroalgal kelp forests to thrive. Currently, sea otters are highly fragmented across predation refuges within islands, where their ecological function has been restricted to very shallow waters (~ 3 m) as compared to deeper adjacent habitats (>4 – 8 m; left column) where they once affected and maintained kelp forests. Photos are from Kiska Island in August 2021, where divers swam a perpendicular line from 8 m depths to 3 m depths documenting the changes in kelp and urchins in a predation refuge occupied by otters (top right). At the 3 m depth, there was evidence of sea otter predation on sea urchins by field-based observers (right, 2^nd^ from top) and in urchin debris on the seafloor (middle right). Despite evidence of predation on sea urchins at 3 m depth, sea urchins were available within crevices in the complex reef habitat, potentially shifting in from areas of high density at > 4 m depths into areas of lower density at 3 m depth following a pattern of source – sink dynamics (bottom right) (Photo Credit: U.S. Fish and Wildlife Service).

An updated estimate of sea otter equilibrium density from the prior 15 otter/km^2^ estimate to 11.8 otters/km^2^ is also informative from a population monitoring perspective. The original estimate of 15 otters/km^2^ was derived based on the average raw otter density from aerial surveys (4.52 otters/km^2^) multiplied by an aerial to skiff correction factor (3.58) divided by the estimated total available habitat to sea otters (6,503 km^2^) (Burn et al. 2003). This correction factor, which was calculated as the ratio of counts made from aerial counts to near simultaneous skiff-based counts in the 1990s and 2000s, was previously broadcast to sea otter counts made across islands and hindcast to surveys dating back to the 1960s (Burn et al. 2003). In our current model we relaxed the assumption of a fixed correction factor and allowed detection probability (inverse of correction factor) to vary by population size and island size. This strategy provided more robust estimates of detection probability and correction factors, and thus an improved estimate of equilibrium density. Despite this, current data limitations required that we use a fixed estimate for skiff-based detection probability at 0.8. We acknowledge detection probability in skiff surveys likely also varies with island size, sea otter population status, and other factors such as coastline complexity, food resources, and predation risk. Recent sea otter skiff survey efforts in the Southwest DPS have attempted to incorporate distance sampling (Buckland 2001, Kéry and Royle 2016), which could provide more precise estimates of detection probability in the future (U.S. Fish and Wildlife Service, unpublished data).

Last, the ability of wildlife managers to mitigate threats sufficiently to ensure persistence of sea otters is complicated by the nature of the threat (killer whales) that has caused sea otter decline and is hindering recovery. Killer whale predation reduced the population at each island leading to an overall population decline in the region. The remaining sea otters have exhibited restricted habitat use resulting in patchily distributed clusters of sea otters within predation refuges that indicate little sign of connectivity. Future research could investigate rates of dispersal between occupied patches and the success of dispersers. We used dispersal probabilities of 0.05 for sub-adults and 0.01 for adults of both sexes, based on previously published estimates of sea otter movement rates over similar spatial scales (Garshelis and Garshelis 1984, Ralls et al. 1996, Tinker et al. 2008, Laidre et al. 2009, Tarjan and Tinker 2016, Breed et al. 2017). We have limited current information on the rate at which sea otters in the region disperse and the success rate of dispersers. Additional VHF or GPS-tagging studies of sea otters or genetic assessments, possibly through collection of biological samples (e.g., tissue, hair), could improve our understanding of dispersal rates and connectivity within islands and across islands (Davis et al. 2019, Conn et al. 2020, Feutry et al. 2020). Information on whether sea otters are attempting to disperse and are unsuccessful due to predation (or other causes of mortality), or that sea otters are averse to dispersal (and dying locally) as a result of either a behavioral or genetic predisposition favoring non-dispersal (Elliot et al. 2014) is an unstudied but crucial component tied to potential management actions. Distinguishing between these scenarios will help guide whether reintroductions are worthwhile as a management action to aid sea otter recovery (Davis et al. 2019). These studies could also provide updated estimates of sea otter survival rates, which could help refine future sea otter viability projection models. This line of inquiry could provide further clarity on the extent to which killer whales are currently influencing vital rates, or whether the observed patterns of sea otter population trends are a relic of killer whales past (Sheriff et al. 2010).

## Acknowledgements

We appreciate the efforts by many researchers involved in understanding sea otters across the Aleutian Archipelago over the past six decades from the U.S. Geological Survey, U.S. Fish and Wildlife Service, University of California Santa Cruz, and many other institutions. This manuscript benefited from internal reviewers at the U.S. Fish and Wildlife Service and the U.S. Geological Survey. This study was supported by the U.S. Fish and Wildlife Service Marine Mammals Management Program and the U.S. Air Force with logistical support provided in 2021 by the crew of the *R/V Norseman II*, in 2015 by the U.S. Fish and Wildlife Service National Wildlife Maritime Refuge *R/V Tiĝlax̂*, and multiple vessels and partners from the National Science Foundation in the preceding and interim years. Most of the sea otter population survey data is publicly available through the Kenner et al. 2021 data release. The 2021 survey is available from the U.S. Fish and Wildlife Service. This manuscript has been approved for publication consistent with USGS Fundamental Science Practices (http://pubs.usgs.gov/circ/1367/). Any use of trade, firm, or product is for descriptive purposes only and does not imply endorsement by the U.S. Government. We offer respect and acknowledgement to the Aleut or Unangax̂ whose lands and waters we surveyed and learned so much from in the Near Islands (*Sasignan Tanangin*) at Attu (*Atan*), Agattu (*Angatux̂*), and Semichi Islands (*Samiyan*); the Rat Islands (*Qax̂ um tanangis*): Kiska (*Qisxa*), *Hawadax* (restored from “Rat” to traditional Aleut name in 2012), and Amchitka (*Amchixtax̂*) and the Adreanof Islands (*Niiĝuĝim tanangis*): Ogliuga and Skagul (Aglaga and Sxaĝulax), Tanaga (Tanax̂ ax), and Adak and Kagalaska (Adaax and Qigalaxsix̂).

---

**Figure S1.**
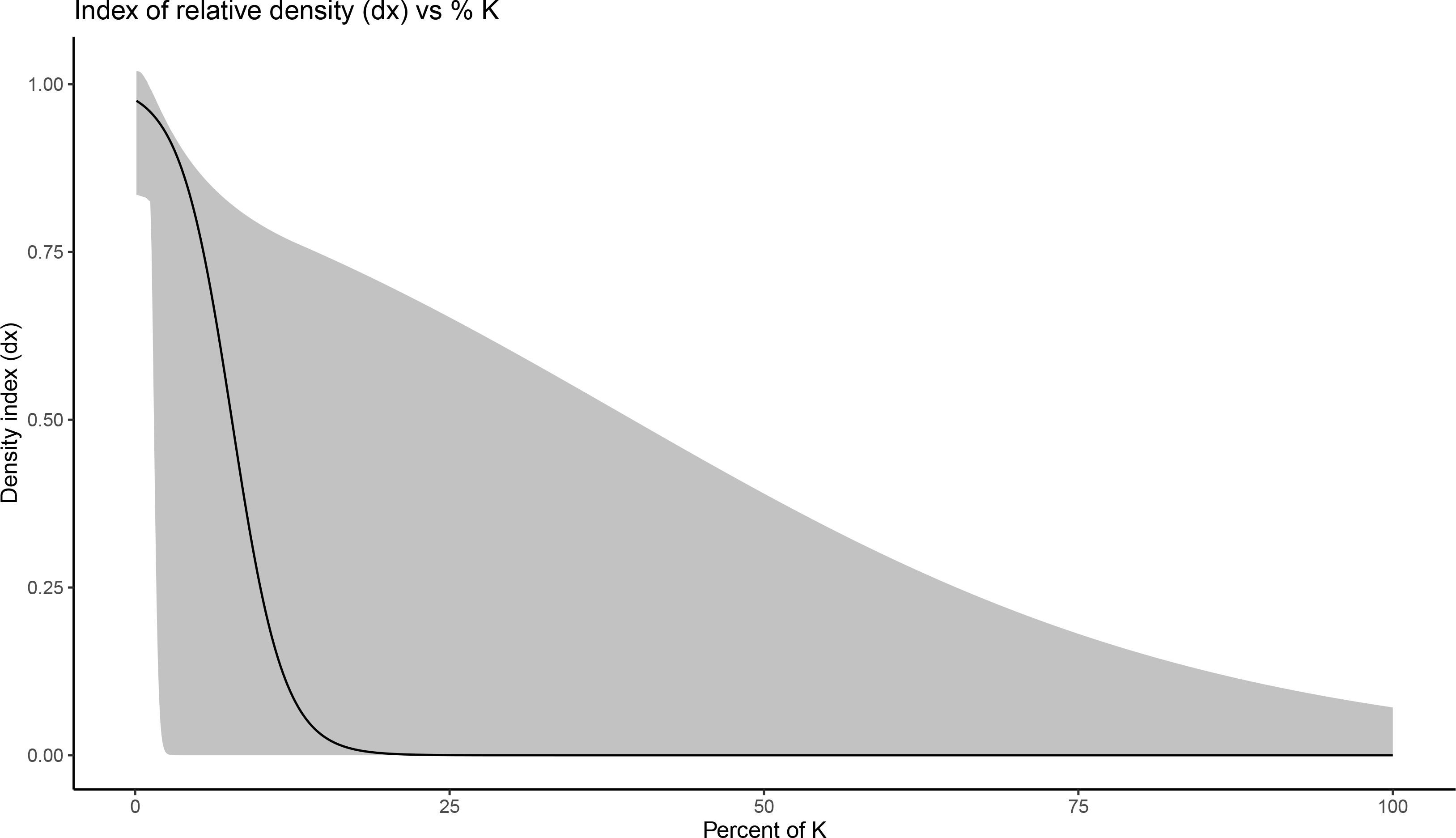

**Figure S2.**
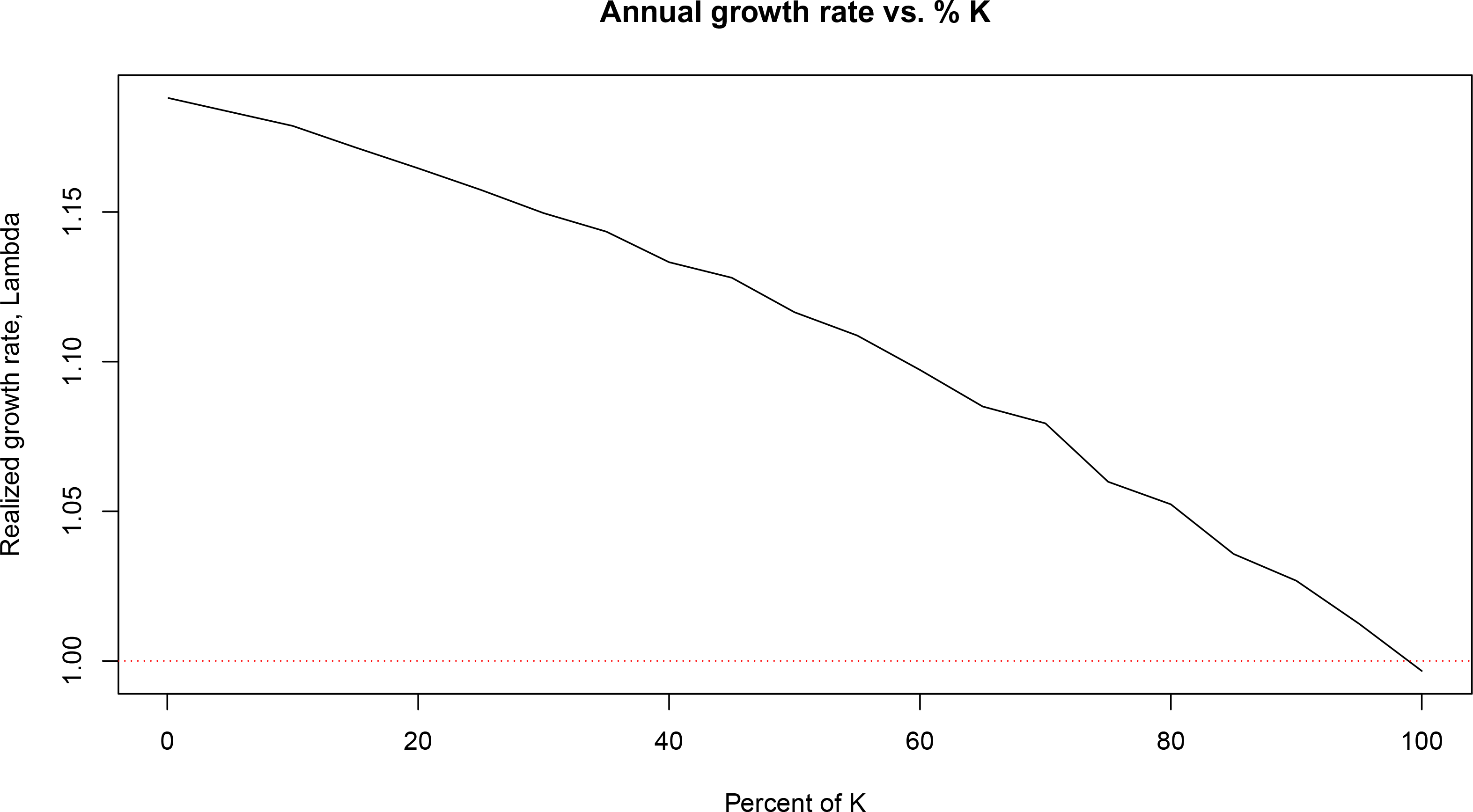

**Figure S3.**
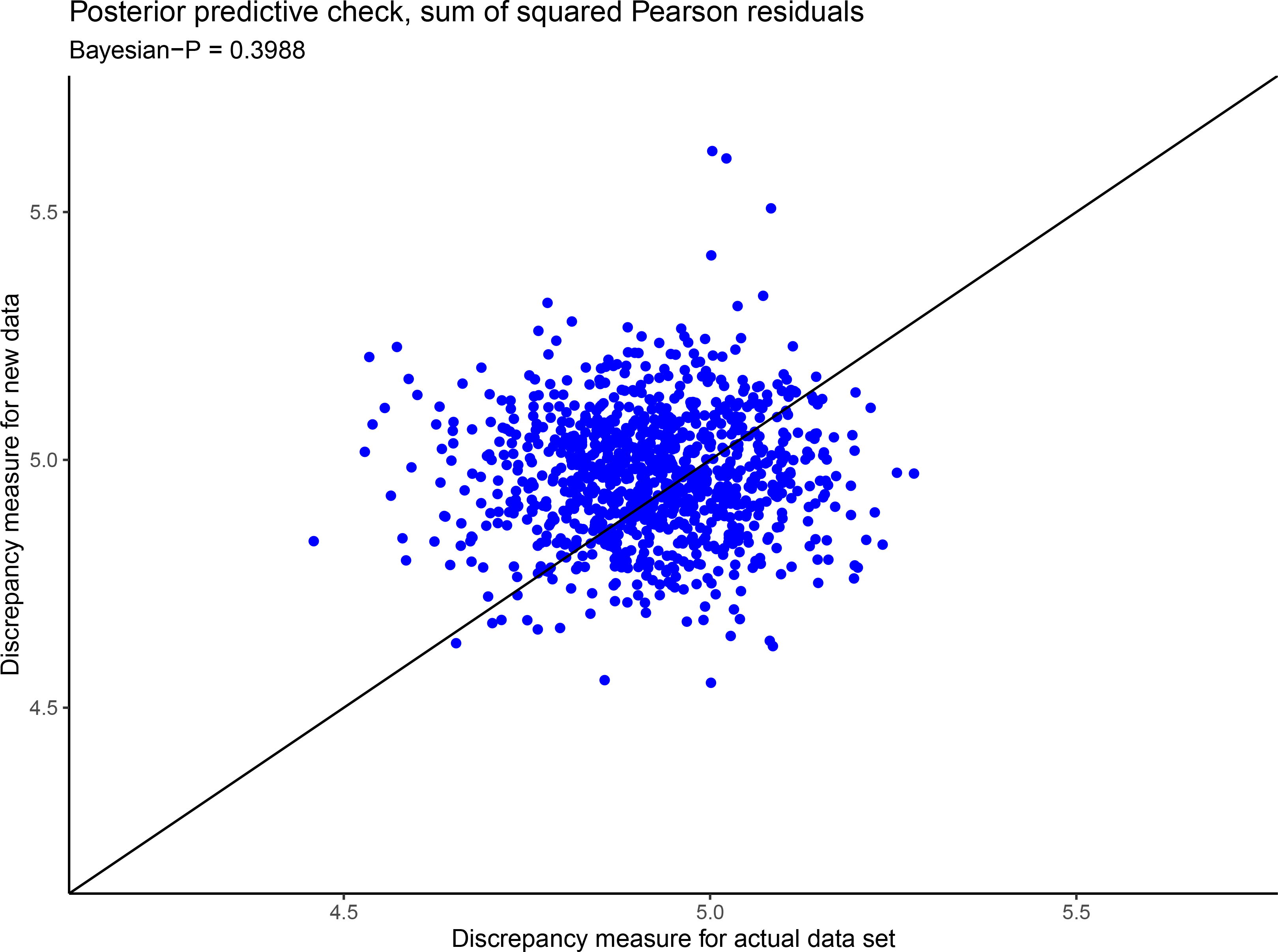

**Figure S4.**
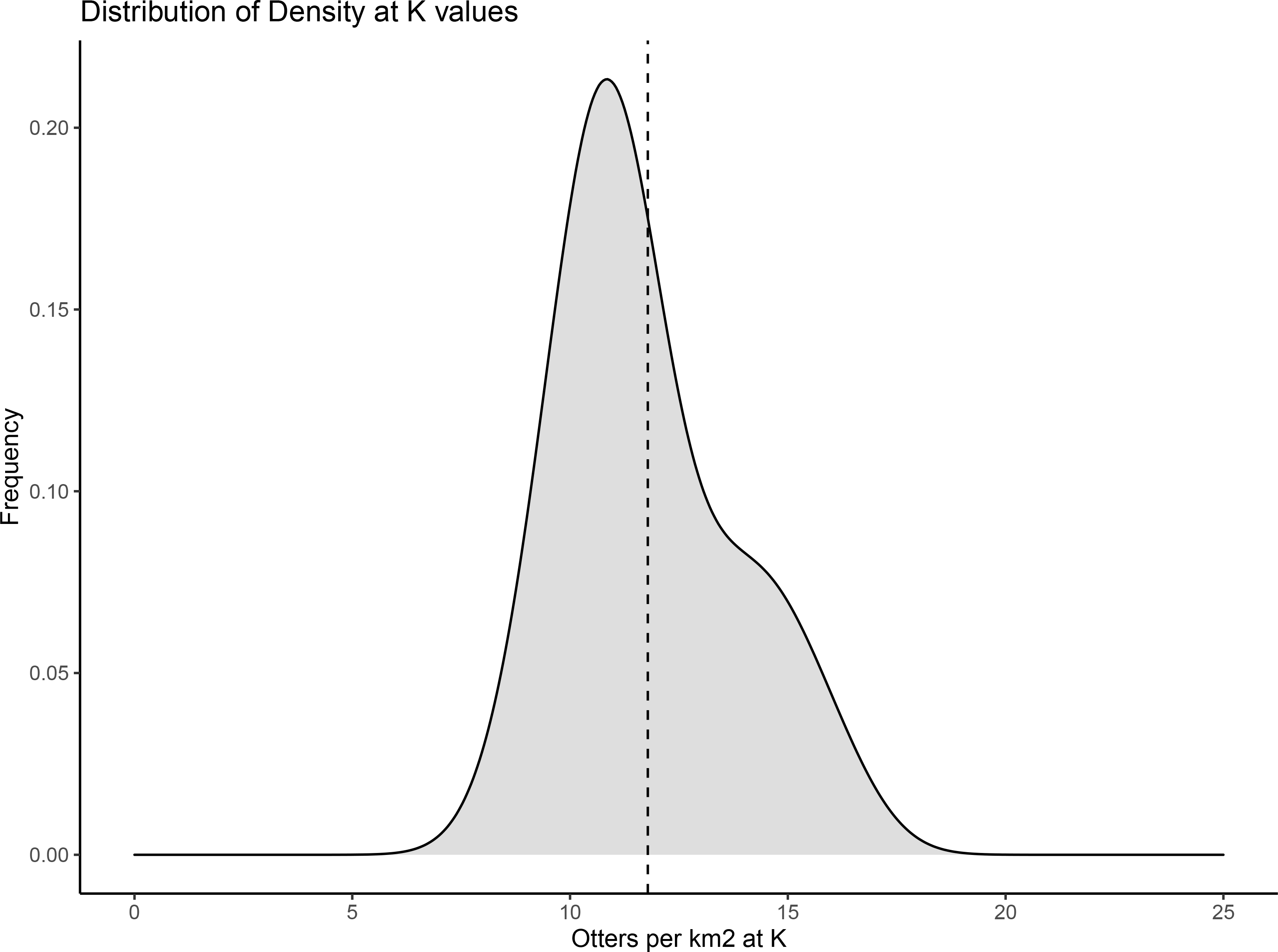

